# Dynamic neuro-immune regulation of psychiatric risk loci in human neurons

**DOI:** 10.1101/2024.07.09.602755

**Authors:** Kayla G. Retallick-Townsley, Seoyeon Lee, Sarah E. Williams, Sam Cartwright, Sophie Cohen, Annabel Sen, Meng Jia, Hannah Young, Lee Dobbyn, Michael Deans, Meilin Fernandez-Garcia, Laura M. Huckins, Kristen J. Brennand

**Affiliations:** Department of Genetics and Genomics, Icahn School of Medicine at Mount Sinai, New York, NY, USA; Icahn Institute of Genomics and Multiscale Biology, Icahn School of Medicine at Mount Sinai, New York, NY, USA; Pamela Sklar Division of Psychiatric Genomics, Icahn School of Medicine at Mount Sinai, New York, NY, USA; Nash Family Department of Neuroscience, Friedman Brain Institute, Icahn School of Medicine at Mount Sinai, New York, NY 10029; Department of Psychiatry, Division of Molecular Psychiatry, Yale University School of Medicine, New Haven, CT 06511; Department of Genetics, Wu Tsai Institute, Yale University School of Medicine, New Haven, CT 06511

**Keywords:** massively parallel reporter assay, human induced pluripotent stem cells, dynamic expression quantitative trait loci, psychiatric genetics, inflammation, immune signaling, fetal brain development

## Abstract

The immune environment influences neurodevelopment and subsequent clinical trajectories for psychiatric outcomes in childhood and adolescence. Yet it remains unclear if the impact of maternal and fetal immune activation varies with distinct polygenic risk profiles. Therefore, here we catalogue genotype and environment (GxE) interactions, contrasting allele-specific regulatory activity between inflammatory cues. We report a cue-specific neuronal massively parallel reporter assay (MPRA) of 152 loci from genome-wide association study (GWAS) linked to ten brain traits/disorders, empirically dissecting the impact of interleukin-6 (IL-6) and interferon-alpha (IFNα) on transcriptional activity. In human induced pluripotent stem cell (hiPSC)-derived glutamatergic neurons, 1,156 active candidate regulatory risk sequences (MPRA-active CRSs) are resolved, including 267 with variant-specific effects (MPRA-emVars) and 61 with variant-by-cytokine interaction effects (interaction MPRA-emVars). Broadly, neuronal immune-mediated regulatory activity is associated with differences in transcription factor binding and chromatin accessibility, the gene targets of which show pleiotropic enrichments for brain, metabolic, and immune disorders. Dynamic genetic regulation mediates neuroimmune effects, informing our understanding of the genomics of psychiatric and neurological traits, mechanisms governing pleiotropy across disorders, and how immune mechanisms mediate genetic risk.

## Introduction

Genome-wide association studies (GWAS) link hundreds of significant loci with risk for psychiatric traits^1–6^ and neurodegenerative disease^7,8^; the overwhelming majority are non-coding variants, common in the population-at-large, and thought to regulate the expression of one or more target genes^9^. Yet, pinpointing specific causal variants is complex, with true signal obscured by linkage disequilibrium patterns, and polygenic risk scores still incapable of reliably predicting individual outcomes^10^. Interactions between risk variants with each other^11,12^ and the environment^13^ may underlie the variable penetrance and expressivity of genetic risk for complex brain disorders.

Neurodevelopment represents a key window during which environmental exposures interact with genetic risk^14^. Maternal immunometabolic stressors (e.g., infection^15^, trauma^16^, immune dysfunction^17^, obesity^18^, and diabetes^19^) are associated with subsequent neuropsychiatric disorder risk in childhood and adolescence. High levels of maternal cytokines (e.g., interleukin-6 (IL-6)^20,21^) cause immune dysregulation in rodent offspring^21–23^, concomitant with changes in neurodevelopment, gene expression, circuit function, and behavior^24–27^. IL-6 likewise impacts gene expression, neurogenesis, and neuronal activity in cultured human neural cells^28–30^. Likewise, fetal cytokines (e.g., type 1 interferons, including IFNα) mediate placental response to infectious agents and can influence neurological and neurodevelopmental trajectories in humans^31^; for example, alterations in brain development are often present in patients with genetic interferonopathies^32^. Yet, what remains unclear is the extent to which cytokine effects vary between individuals with distinct polygenic risk profiles.

Traditional genomic studies assume genetic risk to be static, and so map risk variants without consideration of how regulation of gene expression can change, but examples of cell-type-^33–35^, sex-^36–38^, and developmental stage-^39–42^ specific, stress-^43,44^, inflammation-^45,46^, body mass index-^47^, and drug-^48–50^ dependent genetic regulation of gene expression abound. We hypothesize that the regulatory activity of non-coding risk variants in human neurons is influenced by inflammatory cues, and that resulting cue-specific effects may alter susceptibility for complex brain disorders and diverse neurotypes. Which genetic variants show dynamic immune-responsive regulatory activity in neurons is unknown.

By coupling massively parallel reporter assays (MPRAs)^51,52^ with human induced pluripotent stem cell (hiPSC) models^53–55^, transcriptional activity can be empirically evaluated at scale in live human neurons. Here, we test the hypothesis that immune signaling (specifically IL-6 and IFNα) interact with non-coding regulatory elements by characterizing the dynamic transcriptional activity of 3,668 candidate regulatory sequences (CRSs) prioritized from 152 GWAS loci linked to ten brain traits/disorders. Altogether, we modelled dynamic immune contributions to neurodevelopment that precede symptom onset and disorder etiology by decades.

## Results

### Dynamic neuronal changes in response to immune signaling

Given that hiPSC-derived *NGN2*-induced glutamatergic neurons (iGLUTs)^56,57^ most resemble their fetal counterparts^58^, the influence of inflammatory cues during neurodevelopment was assessed using iGLUTs from two neurotypical donors (one female, one male) that were acutely (24- and 48-hours) treated with IL-6 (25ng/ml, 60ng/mL), IFNα-2b (100 IU/mL, 500 IU/mL), or vehicle (0.1% FBS in ultrapure H_2_O) before harvesting at 24 days *in vitro* (DIV) (experimental schematic: **SI Fig. 1**). Treatment dose was informed by previous studies in human neural progenitor cells (NPCs) and neurons: IL-6^28,30^, IFNα^59^.

Receptors for all stressors (IL-6: *IL6R*, *IL6ST;* IFNα: *IFNΑR1*, *IFNAR2*) were expressed in iGLUTs (**SI Fig. 2**). In classical IL-6 signaling, IL-6 binds to membrane-bound IL6R, which then associates with glycoprotein 130 (GP130, encoded by *IL6ST*); comparatively, trans-signaling via hyper-IL-6 delivers IL-6 covalently bound to soluble IL6R. Notably, no qualitative differences between classical and trans IL-6 signaling pathways have been reported^60^. Whereas some reports indicate that hiPSC-derived NPCs express low *IL6R,* do not respond to IL-6, and require hyper-IL-6 treatment^28,61^, others find that IL6-R mRNA and protein is expressed in hiPSC-derived neurons^62^, increases with maturation^62^, and that IL-6 treatment impacts expression of genes regulating extracellular matrix, actin cytoskeleton and TGF-beta signaling^63^. Here, we likewise demonstrate that DIV23 iGLUTs express *IL6R* (**SI Fig. 2A-C**), albeit at relatively low levels (0.5-1 normalized TPM), and demonstrate neuronal transcriptomic, epigenomic, and cellular response to classical IL-6 signaling as follows.

In glia, different immune stressors induce distinct cellular states^64,65^; our findings indicate that this may also occur in neurons (**SI Figs. 3-6**). There was limited overlap between either significant (BH-FDR-corrected p-value p_FDR_<0.05) or nominally significant (unadjusted p_nom_<0.05) differentially expressed genes (DEGs) by cue, with IL-6 treatment more moderately impacting the transcriptome (12 down-regulated and 1 up-regulated DEG, p_FDR_ <0.05: 1434 DEGs, p_nom_<0.05) relative to IFNα treatment (39 down- and 97 up-regulated DEGs, p_FDR_ <0.05; 2000 DEGs, p_nom_<0.05) (**SI Fig. 3A-B; SI Data 1.1-1.4**). Individual cytokine significant effects at 24 and 48 hours were significantly correlated (IL-6: r=0.35, p=0.003; IFNα r=0.72, p=5.7×10^-230^) (**SI Fig. 3A-C**), albeit with differences in the magnitude of effects over time. Nominal IL-6 DEGs were enriched for processes largely related to cell adhesion, whereas IFNα response genes were enriched for antigen binding, protein ubiquitination, and as expected, interferon (-log(p)=9.02; z-score=2.287) and neuroinflammatory signaling (-log(p)=8.24) (**SI Fig. 3D-E; SI Data 1.5**).

Although neuronal immune responses were distinct between exposures, convergent mechanisms and shared enrichments between IL-6 and IFNα response were also resolved. Both treatments yielded DEGs (p_nom_<0.05) enriched for pathways related to stress and immune response (e.g. mTOR signaling [IL-6(-log(p)=3.7; IFNα (-log(p)=6.25] and EIF2 signaling [IL-6(-log(p)=10.2; IFNα (-log(p)=24.7)]) (**SI Data 1.5**) and well-recapitulated fetal mouse brain signatures associated with four models of immune activation (poly(I|C)^66^ (**SI Data 1.6)**. These similarities likely reflect a set of convergent genes (258 down-regulated and 352 up-regulated) with perturbations in the same direction across neuroinflammatory contexts (meta-analysis p_FDR_<=0.05) (**SI Data 1.7**).

Chromatin accessibility changes were greatest with IL-6 (675 differentially active regions (DARs); p_FDR_ <=0.05) and more modest with IFNα (12 DARs; p_FDR_ <=0.05) (**SI Fig. 4-5; SI Data 1.9-1.11**), unlike transcriptomic changes, which were greatest with IFNα. Yet, of those DEGs that overlapped with chromatin DARs, cue-responsive gene expression (average expression) and chromatin accessibility (ATAC peak score) were significantly, albeit weakly, positively correlated [IL-6 exposure: Pearson’s Correlation Coefficient r=0.17, p=0.001; IFNα exposure: r=0.13, p=0.0004)] (**SI Fig. 5A**). Comparative enrichment analysis revealed robust IL-6-specific enrichments for calcium-dependent signaling and O-linked glycosylation, IFNα-specific enrichments for interferon signaling and synaptic transmission, and shared enrichments for actin/cadherin binding (**SI Fig. 5B-C**).

Phenotypically, neither cytokine altered cell number or synaptic puncta density, but IL-6 increased the number of mature neurons (p_bon_<0.01) and IFNα significantly increased neurite outgrowth (p_bon_<0.001) in immature neurons (**SI Fig. 6**).

Altogether, multimodal evidence indicated that acute exposure to IL-6 and IFNα in mature iGLUTs resulted in distinct neuronal responses.

### Dynamic transcriptional regulation of allele-specific activity at GWAS loci in neurons

To test the extent that neuronal immune response altered genetic regulation of expression by disease-relevant loci, we designed a cross-disorder Lenti-MPRA^52^ library integrating GWAS summary statistics from ten brain disorders, neurotypes, and traits (Alzheimer’s disease (AD)^67^, attention deficit hyper-activity disorder (ADHD)^68^, anorexia nervosa (AN)^69^, autism spectrum disorder (ASD)^3^, bipolar disorder (BIP)^70^, major depressive disorder (MDD)^71^, obsessive compulsive disorder (OCD)^72^, post-traumatic stress disorder (PTSD)^73^, schizophrenia (SCZ)^1^, and neuroticism (NEU-P)^74^) with post-mortem brain eQTLs^75^ (coloc2^76^ and S-PrediXcan^77^) (**SI Fig. 7; SI Data 2.1-2.7**). We prioritized 4,430 GWAS single nucleotide polymorphisms (SNPs), with the number selected per trait dependent on the number of significant loci per GWAS (**SI Fig. 7C**). The following benchmark variants were included: 310 empirically active MPRA-CRS [164 positive (active with variant-specific) and 146 negative (active without variant-specific effects) controls^78^, and 88 coloc2-controls (lack of colocalization or brain QTL regulation (pph3>0.9)). In total, we included 8,960 sequences (8,860 SNP-centered elements, 100 scramble controls to measure basal activity of the minimal promoter), representing 4,430 biallelic pairs (**SI Data 2.8-2.9**).

The library was transduced into mature iGLUTs (21 DIV); 24 hours later, neurons were acutely exposed (48 hours) to IL-6 (60ng/mL), IFNα (500 IU/mL) or vehicle (vehicle: 0.1% FBS in ultrapure H_2_O) before harvest (24 DIV) (two control donors, two biological replicates each, experimental schematic: **SI Fig. 1**). Technical replicates were highly correlated based on DNA and RNA counts (Pearson’s correlation coefficient=0.98-1.00), log2 (RNA/DNA) ratios across replicates (r=0.61-0.93, mean r=0.83), and mean log2 (RNA/DNA) between donors (r=0.87-0.95) (**Fig. 1A-C, SI Fig. 8, SI Data 2.11**). Following filtering, 3,440 (baseline), 3,322 (IL-6), and 3,366 (IFNα) CRSs and 44-48 scramble sequences were resolved (3,071 shared sequences, 843 bi-allelic variants, 45 shared scrambles, mean n barcode/sequence=45) (**SI Fig. 8D; SI Data 2.11)**.

Across all experiments, scramble sequences were significantly less active than experimental positive benchmark CRS^78^ (Student’s t-test; scramble-v-positive: baseline p=0.00095, IL-6 p=0.0028, IFNα p=0.00044) or prioritized CRS (Student’s t-test; scramble-v-prioritized CRSs: baseline p=0.0012, IL-6 p=0.0026, IFNα p=0.00092) (**SI Fig. 11B).** Analysis was repeated across three standard pipelines: MPRAnalyze^79^, mpralm^80^, and DEseq2^81,82^: CRS activity, variant-specific differential activity, and cue-by-variant differentially activity was highly correlated between mpralm and DEseq2, but DEseq2 results were affected by p-value inflation (**SI Fig. 9-10**).

**MPRA-CRSs** were defined as “MPRA-validated cis-regulatory sequences” and measured as CRS with either significantly (mpralm p_FDR_<=0.1) higher transcriptional activity (active) or lower transcriptional activity (repressed) compared to the mean transcriptional activity. 26-41% of MPRA sequences were transcriptionally active (baseline, n=1,057 out of 3,440, 31%; IL-6, n=1,092 of 3322, 33%; IFNα, n=764 of 3,366, 22.7%) (**Fig. 1D**); in total, 1,156 CRS were active in at least one context and 660 were active across all contexts (**SI Fig. 12**).

**Figure 1.**
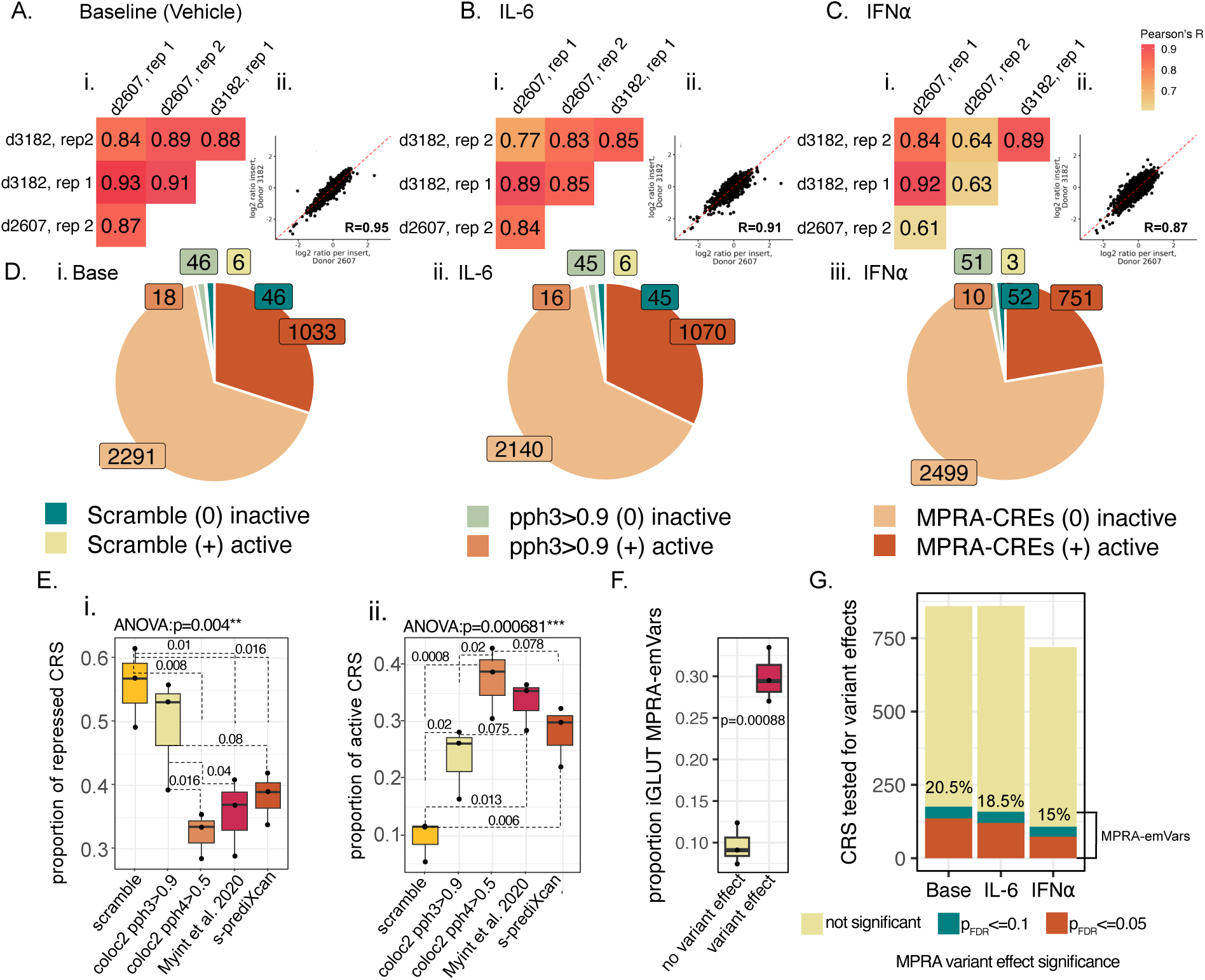
Dynamic immune-responsive neuronal MPRA-emVars, related to SI Fig. 8-13, SI Data 2.1-2.16. Mature hiPSC-derived glutamatergic neurons (iGLUTs; 2 neurotypical donors/2 replicates) transduced with cross-disorder/trait lenti-MPRA at baseline **(A)** (UltraClear-H2O or 0.1% FBS) or when treated with **(B)** 60ng/mL IL-6 or **(C)** 500 IU/mL IFNα-2B, for 48hrs. **(i)** Top left: Following QC and filtering, MPRA activity was strongly correlated (Pearson’s Correlation Coefficient) across biological (donor 1 – donor 2) and technical replicates (sequencing batch rep.1 – rep. 2) within each condition. **(ii)** Top right: mean transcriptional activity across technical replicates was strongly correlated between donors. **(D)** Active MPRA-CRSs were identified as those with a transcriptional rate significantly (p_FDR_<0.1) greater than the average activity (representing the baseline rate of the minimal promoter). 26-31% of MPRA sequences were transcriptionally active (baseline, n=1,057 out of 3,440, 31%; IL-6, n=1,092 of 3,322, 33%; IFNα, n=764 of 3,366, 22.7%). **(i-iii)** Pie charts representing the number of active CRSs by condition. Proportion of active/inactive MPRA-CRS (dark orange/tan) is greater than for scramble (teal/pale yellow) and coloc2-controls (pph3>0.9 salmon/light green). **(E)** The mean proportion of repressed **(E,i)** and active **(E,ii)** sequences significantly differed by prioritization method (repressed ANOVA p=0.004, active ANOVA p=0.000681). Across all experiments, the proportion of active scramble sequences was significantly less than experimental benchmark CRS^78^ (n=3; post-hoc paired Student’s T-tests: scramble-v-positive p=0.013) or variants prioritized by colocalization (pph4>0.5) (n=3; scramble-v-coloc2 p=0.0008) and S-PrediXcan (n=3; scramble-v-prediXcan p=0.006). The proportion of active MPRA-CRSs was significantly greater for eQTL-colocalized GWAS variants (pph4>=0.5) than non-colocalized controls (pph3>0.9) (n=3; post-hoc paired Student’s T-test: p=0.02). **(F)** The proportion of experimental positive benchmark variants^78^ identified as MPRA-emVars (bi-allelic sequences with significant ref-v-alt difference in activity) was significantly greater (paired Student’s T-test, p-value=0.00088) than for negative benchmark variants^78^. Boxplots with lower and upper hinges (the 25th and 75th percentiles), lower and upper whisker that extends from the hinge to the largest or smallest value no further than 1.5 * IQR (inter-quartile range). Individual samples (unique MPRA experiments n=3) are represented by individual points. Differences in the proportions was evaluated by two-way ANOVA followed by paired two-sided T-tests following testing for assumptions of normalcy (Shapiro-Wilk’s Test) and equal variance (Levene’s Test). **(G)** Across contexts, 15-20.5% of tested variants were MPRA-emVars (teal p_FDR_<=0.1 threshold; terracotta; p_FDR_<=0.05 threshold), with the greatest number of active MPRA-emVars at baseline **(SI Fig. 13C)**.

The mean proportion of repressed **(Fig 1E,i)** and active **(Fig 1E,ii)** sequences was significantly different between scramble sequences, positive reference sequences^78^, and prioritized CRS (active ANOVA p=0.000681, post-hoc pairwise Student’s T-test, n=3: scramble-v-positive p=0.013, scramble-v-coloc2 controls p=0.02, scramble-v-coloc2 p=0.0008, scramble-v-PrediXcan p=0.006) (**SI Fig. 12B**). Notably, relative to coloc2-controls (pph3>0.9), top eQTL-colocalized GWAS sequences (pph4>=0.5) had a significantly greater proportion of active MPRA-CRSs (n=3; post-hoc paired Student’s T-test: coloc2-controls (pph3>0.9)-v-eQTL-colocalized GWAS p=0.02) (**Fig. 1E,ii**). Moreover, CRS prioritized by colocalization (coloc2 pph4>0.5) were more likely to be active CRS compared to CRS prioritized by transcriptomic imputation (coloc2-v-PrediXcan p=0.078).

**MPRA-emVars** were defined as “MPRA-active CRSs with single-nucleotide (variant) changes that significantly altered transcriptional activity” and are calculated as variants with a significant difference in transcriptional activity between the reference allele and the alternative allele (mpralm variant-specific, p_FDR_<=0.1), restricted to those with positive CRS activity (mpralm quantification p_FDR_<=0.1, logFC>0). Of the 859 (base), 860 (IL-6), and 719 (IFNα) transcriptionally active biallelic MPRA sequences (**SI Fig. 13A-B**), 15-20% (20.5% baseline, n=176; 18.5% IL-6, n=159; 15% IFNα, n=108) showed variant specific effects (BH-FDR-corrected p-value p_FDR_<0.1) **(Fig. 1G; SI Data 2.12-14**). The proportion of positive benchmark variants (selected based on empirical differential variant activity^78^) that were MPRA-emVars was greater than the proportion of negative benchmark variants^78^ (n=3, paired Student’s T-test, p=0.00088) (**Fig. 1F**). Allelic shifts were significantly correlated across cues (no significance threshold: r=0.85-0.88, p<2.2×10^16^, p_FDR_<0.1: r=0.96-0.98, p<2.2×10^16^; **SI Fig. 13C**) and replicate those reported in human neural progenitor cells^54^ (MPRA-emVar p_FDR_<0.1 threshold at baseline: r=0.9, p=0.0055) (**SI Data 2.17**), although there are too few overlapping CRS (n=76) for this to represent robust validation (**SI Data 2.18**).

### Concordance of iGLUT MPRA-emVars with eQTL effects in the adult and fetal brain

Effect sizes of concordant MPRA-emVars were significantly correlated with eQTL betas (fetal cortex Pearson’s R=0.66-0.69; excitatory neurons R=0.56-0.83; adult pre-frontal cortex R=0.58-0.6) (**SI Fig. 14D**). Of course, MPRAs are synthetic measures of transcriptional activity outside the endogenous context, whereas eQTLs measure variant regulatory activity in the native genome, where one SNP can regulate multiple genes (eGenes) with different magnitudes and directions of effect. In particular, human fetal brain has more SNPs that map to multiple eGenes with opposing directions of regulation (**SI Fig. 14A-C**). To represent this nuance, we considered overlap of MPRA-emVars based on the absolute number of regulatory SNPs (**Fig. 2A**) and based on the eQTL effects (SNP-eGene associations) (**Fig. 2B**). At baseline, 22%, 29%, and 22% percent of MPRA-emVars overlapped with significant (Bonferroni or FDR<=0.05) eQTLs in the adult cortex^75^, fetal brain^83^, or adult cortical excitatory neuronal^84^, the majority of which were concordant in direction with at least one SNP-eGene pair in adult cortex (69%), fetal brain^83^ (71%), and excitatory neurons^84^ (55%) (**Fig. 2A**). Concordance when considering all SNP-eGene pairs was highest in excitatory neurons^84^ (68.8%) (**Fig. 2B**), regardless of significance threshold (n=3, paired Student’s t-test; all variants p=0.062, MPRA-emVars p=0.084). Whereas strong concordance with adult cortical eQTLs was only observed when sub-setting for significant MPRA-emVars (n=3, paired Student’s t-test p=0.035) (**Fig. 2B, i-iii).** Overall, MPRA effects were concordant with eQTL data, except between response to IFNa and fetal PFC eQTLs (**Fig. 2B, iv-vi**). MPRA-emVars that overlapped with eQTLs (SNP-eGene pairs) identified by S-PrediXcan and coloc2 were highlighted (**Fig. 2C**).

**Figure 2.**
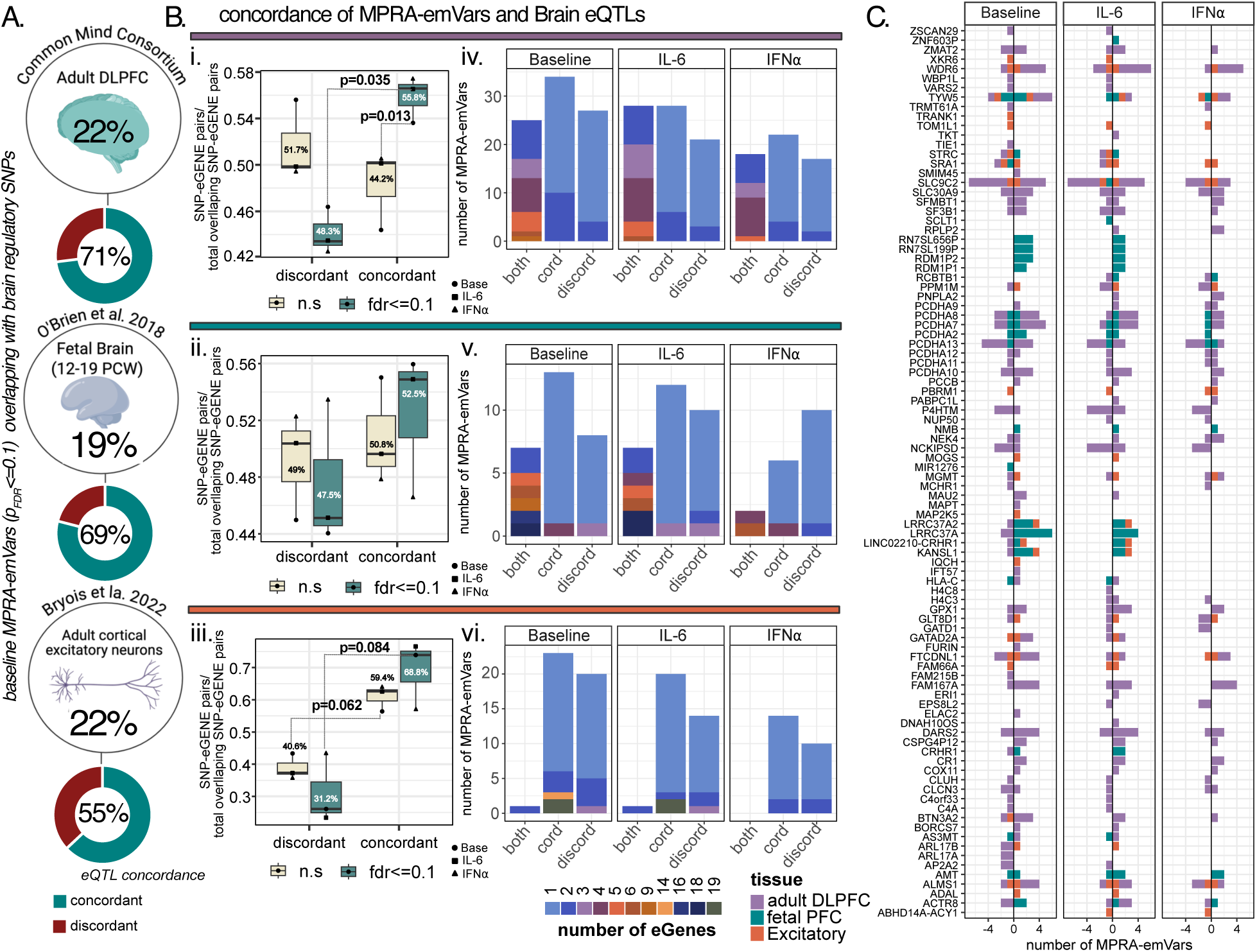
Neuronal MPRA-emVars largely validate brain eQTLs across inflammatory exposures, related to SI Fig. 15. **(A)** 22%, 29%, and 22% percent of MPRA-emVars at baseline overlapped with significant (Bonferroni or FDR<=0.05) **(i)** adult brain^75^, **(ii)** fetal brain (12-19 PCW^83^), and **(iii)** adult cortical excitatory neuronal^84^ regulatory SNPs, respectively. Of these, 69%, 71%, and 55% had concordant directions of effect. **(B)** The proportion of MPRA-emVars with concordant directions of effect with **(i)** adult DLPFC, (**ii**) fetal brain, and **(iii)** excitatory eQTLs (SNP-eGene pairs). Boxplots with lower and upper hinges (the 25th and 75th percentiles), lower and upper whisker that extends from the hinge to the largest or smallest value no further than 1.5 * IQR (inter-quartile range). Individual samples (unique MPRA experiments n=3) are represented by individual points. Average percent concordance is annotated within the boxes, differences in the proportions was evaluate between pairs using two-sided Student’s T Tests after testing for assumptions of normalcy (Shapiro-Wilk’s Test) and equal variance (Levene’s Test). The absolute number of MPRA-emVars that mapped to SNPs that regulated multiple eGenes with concordant (same direction between MPRA logFC and eQTL beta), discordant (opposing direction between MPRA logFC and eQTL beta), or both effects (up and down regulation dependent upon the eGene) in **(iv)** adult DLPFC, (**v**) fetal brain, and **(vi)** excitatory neurons. **(C)** Concordance of MPRA-emVars with their eQTL effects after filtering for S-PrediXcan and coloc2 prioritized eGenes at **(i)** baseline or **(ii)** after IL-6 or **(iii)** IFNα exposure. Bars show the frequency of MPRA-emVars mapping to an eGene across tissues (purple=adult DLPFC, teal=adult Excitatory neuron, orange=Fetal PFC). The x axis represents the total number of MPRA-emVars with the same direction of effect (positive values) or opposite direction (negative values).

### Neuronal immune activation results in significant cue-by-variant interaction effects on transcriptional activity enriched for psychiatric traits and common co-morbidities

152 brain trait GWAS loci were represented by 843 biallelic variants measured in baseline, IL-6, and IFNα conditions; 52 GWAS loci had variant effects across all three conditions, and 22 GWAS loci were specific to one immune condition (14 IL-6, 6 IFNα) (**Fig. 3A,B**). MPRA-emVars overlapped with significant variants in multi-trait or pleiotropic GWAS and highlight differential regulation of genetic risk by neuronal immune activation (e.g., IL-6 MPRA-emVars were annotated for SCZ-BIP pleiotropy and IFNα MPRA-emVars were annotated for Parkinson’s disease) (**Fig. 3C, SI Fig. 15, SI Data 2.19**).

**Figure 3.**
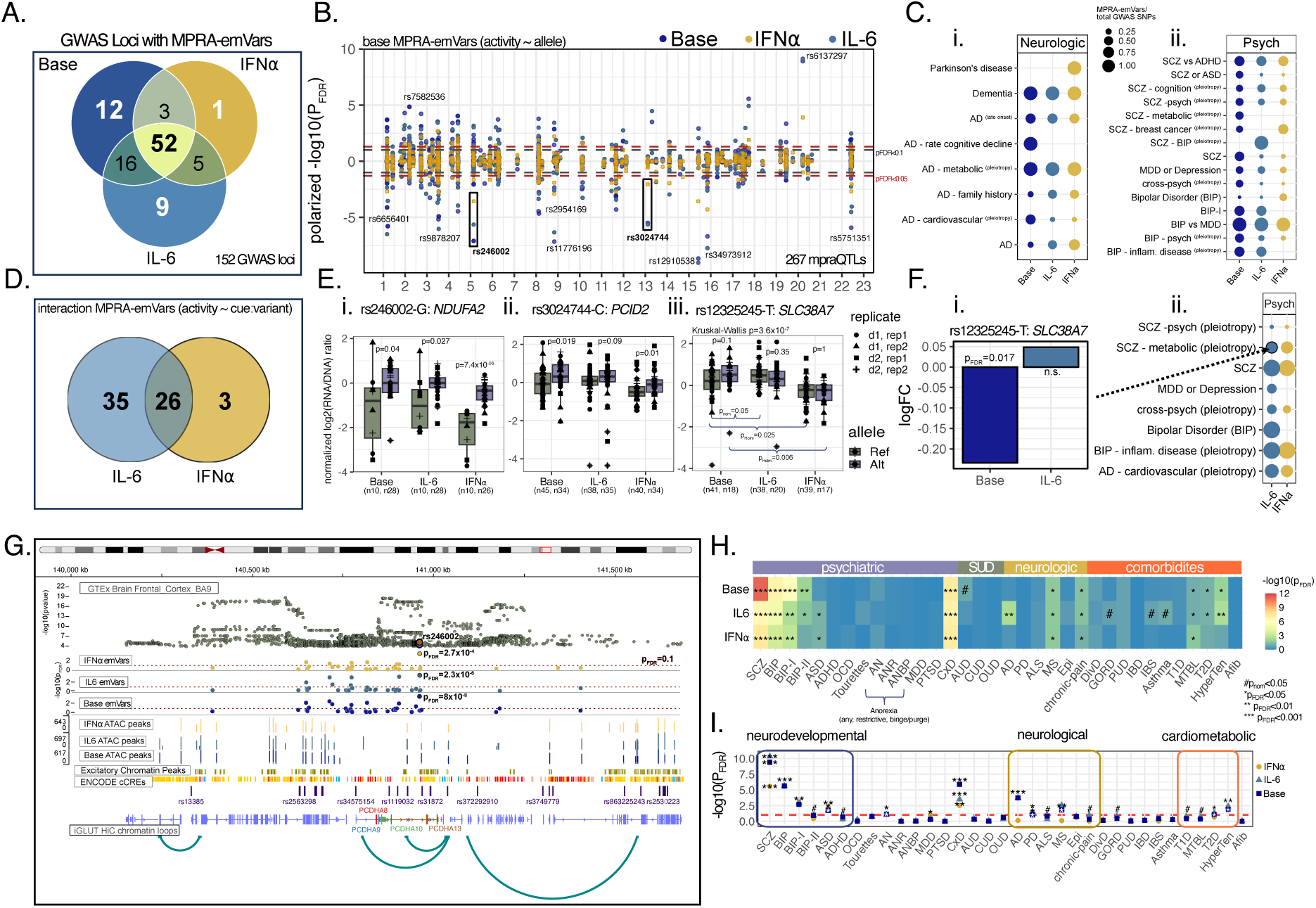
Neuronal inflammation alters the effect of psychiatric risk variants on transcriptional activity, related to SI Fig. 14-16, SI Data 2.12-2.16, 2.19. **(A)** While the majority (98 out of 152) of GWAS loci represented had at least one MPRA-emVar at baseline, MPRA-emVars uniquely mapped to 14 GWAS loci for IL-6, and 6 loci for IFNα that did not have a significant hit at baseline**. (B**) 267 active MPRA-CRS had variant-specific effects QTLs in at least one condition. Manhattan plot of MPRA-emVar across conditions represented as the polarized (direction of effect of the ref-alt allele) - log10(p_FDR_). **(C)** Many of these MPRA-emVars are significant GWAS SNPs for psych-psych pleiotropy or psych-cardiometabolic pleiotropy (e.g., SCZ-metabolic pleiotropy and Alzheimer’s Disease (AD) and cardiovascular pleiotropy). Size of the points represents the proportion of MPRA-emVar compared to non MPRA-emVars that are annotated for the same GWAS trait. **(D)** Of active MPRA CRS, 64 (IL-6, n=61; IFNα, n=29) had a significant cue-variant interaction. **(E) (i)** Representative stable MPRA-emVar: rs246002 on chromosome 5, with significantly decreased reference allele activity (grey) across all conditions (within condition Wilcoxon Rank Sum test (ref-vs-alt): base p=0.04, IL-6 p=0.027, IFNα p=7.4×10^-6^). (**ii**) Representative dynamic MPRA-emVar: rs3024744 on chr 13 with significantly decreased reference allele activity (grey) transcriptionally repressed following IFNα exposure (mpralm quant, p_FDR_=0.004, logFC=-0.33) but not IL-6. **(iii)** Representative specific MPRA-emVar: rs12325245 on chr 22 with significantly decreased reference allele activity (grey) only at baseline (baseline ref-v-alt Wilcoxon Rank Sum test p=0.1; interaction test Kruskal-Wallis p=3.6×10^-7^, post-hoc Dunn’s test base-v-IFNα ref p=0.025, base-v-IFNα alt p=0.006). Boxplots with lower and upper hinges (the 25th and 75th percentiles), lower and upper whisker that extends from the hinge to the largest or smallest value no further than 1.5 * IQR (inter-quartile range). Individual points represent unique barcodes, with shape of the points corresponding with biological and technical replicates. **(F) (i)** rs12325245 (risk allele T; *SLC38A7*) is a GWAS variant associated with SCZ-metabolic syndrome pleiotropy; the non-risk allele had greater regulatory activity only at baseline (mpralm p_FDR_=0.017; interaction mpralm (IL-6-v-Base p_FDR_=0.025; not an active IFNα CRS). **(ii)** MPRA-emVars included GWAS variants annotated for psych-psych pleiotropy or psych-cardiometabolic pleiotropy, schizophrenia (SCZ), major depressive disorder (MDD), and Alzheimer’s Disease (AD). **(G-I)** Activity-by-contact (ABC) enhancer-gene interaction scoring: **(G)**Track annotation of the stable emVar, rs246002, which regulates transcription across conditions at the PCDHA locus. Annotations with cue-dependent chromatin accessibility peaks and iGLUT-specific chromatin loops represent the importance of integrating context and cell-type specific multiomic information to add endogenous context to MPRA-emVars. **(H)** Across conditions, MPRA-emVar gene targets were strongly enrichened for SCZ, BIP, and cross-disorder pleiotropy, and common psych-comorbidities like T2D, hypertension, and GORD. Heatmap of MAGMA enrichments across psychiatric, substance use (SUD), neurological, metabolic/autoimmune disorders, and cardiometabolic GWAS, colored by –log10(p_FDR_) (p_nom_<=0.05^#^, p_FDR_ <0.05*, p_FDR_ <0.01**, p_FDR_ <0.001***). **(I)** ABC gene targets of interaction MPRA-emVars revealed cue-responsive regulation by IL-6 and IFNα affected genes involved in neurodevelopment, neurological disorder, and cardio-metabolic traits. Each point represents a gene list of either **(i)** IFNα interaction MPRA-emVar gene targets or **(ii)** IL-6 interaction MPRA-emVar gene targets. Color of the point indicates the MPRQ-QTL values used to calculate the ABC score; red dashed line represents a P_FDR_<0.1 significance cut-off. **Abbreviations:** *<u>NeuroPsych</u>*: Attention Deficit/Hyperactivity Disorder (ADHD), Anorexia (AN; R - restrictive or BP – binge/purge), Autism Spectrum Disorder (ASD), Bipolar Disorder (BIP), BIP type-1 (BIP-I), BIP type-2 (BIP-II), Cross 8 neuro-psych disorders (CxD), Major Depressive Disorder (MDD), Post-traumatic stress disorder (PTSD) , Obsessive-compulsive Disorder (OCD); *<u>Substance Use Disorders:</u>* Alcohol Use Disorder (AUD); Cannabis Use Disorder (CUD), Opioid Use Disorder (OUD); *<u>Neurological:</u>*Alzheimer’s Disease (AD), Parkinson’s Disease (PD), Amyotrophic lateral sclerosis (ALS), Multiple Sclerosis (MS), Epilepsy (Epi); *<u>Inflammatory-Gastrointestinal</u>*: Diverticular Disease (DivD), Gastro-esophageal reflux disease (GORD), Peptic Ulcer Disease (PUD), Inflammatory Bowel Disease (IBD), Irritable Bowel Syndrome (IBS), Asthma, Type-I Diabetes (T1D); *<u>Cardiometabolic</u>*: Metabolic Syndrome (MTBL), Type-2 Diabetes (T2D), Atrial Fibrillation (Afib), hypertension (HyperTen); *<u>Anthropometric:</u>*Body Mass Index (BMI), left-handedness (left hand).

**Interaction MPRA-emVars** were defined as “single-nucleotide changes that significantly altered transcriptional activity in a differential manner between vehicle and either IL-6 or IFNα (mpralm cue*variant interaction test; p_FDR_<=0.1). 18% and 10% of MPRA active CRS resolved significant cue-by-variant interaction effects (IL-6-v-base: 61 of 337, IFNα-v-base=29 of 287; q_storey_<0.1) (**Fig. 3D**). Across cues, 267 unique variants showed significant regulatory effects in at least one condition (**Fig. 3B**), with 64 showing significant cue-by-variant interaction effects (**Fig. 3D**).

Whereas some top loci were stable across all conditions (e.g., rs246002 [**Fig 3Ei**: baseline p=0.04, IL-6 p=0.027, IFNα p=7.4×10^-6^, Wilcoxon Rank Sum Test ref-vs-alt allele]), others were dynamically regulated by immune activation (**Fig 3D-G)** (e.g., rs3024744 [**Fig3Eii**: consistent allelic effects, but variable magnitudes of effect] and rs12325245 [**Fig. 3Eiii**: significant variant effects at baseline only (Kruskal-Wallis: p=3.6×10^-7^; Post-hoc Dunn’s test: base-ref vs. IL-6-ref p_nom_=0.05, base-ref vs. IFNα-ref p_Holm_=0.025, base-alt vs. IFNα-alt p_Holm_=0.006)]. 44-52% of the 64 interaction MPRA-emVars were immune-specific: active CRSs with a significant variant interaction effect only at baseline or only after with IFNα and/or IL-6 exposure (**Fig. 3D**). For example, the SCZ-metabolic pleiotropy risk variant (rs12325245-T; *SLC38A7* [**Fig. 3Eii**, **Fig. 3F]**) had significantly lower activity than the non-risk allele at baseline (p_FDR_ =0.017), showed no allelic effect following IL-6 exposure despite being a highly active MPRA-CRS (mpralm interaction p_FDR_=0.02), but was transcriptionally repressed following IFNα exposure.

Conversely, 38-48% of interaction MPRA-emVars were “immune-responsive”, significant emVars across cues but with significant IFNα and/or IL-6 interaction effects, albeit with the direction of effect retained across cues. Notably, no interaction MPRA-emVars were significant QTLs at both baseline and after cue exposure, but with reversed direction of effect across cues **(SI Data 2.15-2.16)**.

Target genes regulated by cue-specific MPRA-emVars were predicted using an adapted activity-by-contact^85^ model (STARE^86^) that incorporated enhancer activity (MPRA activity and donor- and cue-matched neuronal chromatin accessibility), contact (Hi-C contact frequency), and predicted TF binding affinities. Enhancer-to-gene mapping generally replicated the original eGene distance-based associations while also identifying more distal targets. For example, while rs246002 stably regulates transcription across conditions, incorporation of endogenous information at the PCDHA locus show differential chromatin accessibility peaks across conditions and iGLUT-specific chromatin loops, integration of which allows for more specific mapping of emVars to gene targets (**Fig 3G**). MPRA-emVar ABC gene targets were highly enriched for psychiatric disorder risk genes, including SCZ, BIP, and cross neuropsychiatric disorder pleiotropy (CxD) (MAGMA^87^), as well as common comorbid metabolic and immune syndromes (**Fig. 3H**). Notably, when specifically considering interaction MPRA-emVar ABC gene targets (**SI Data 2.21-2.22**), enrichments were stronger for neurodevelopmental and neurological disorders, and non-brain disorders such as type-II diabetes and hypertension **(Fig. 3I).**

### Variant-specific disruptions in transcription factor binding affinities are associated with dynamic inflammatory-responsive regulation of psychiatric risk loci

CRS can regulate gene expression through binding of sequence-specific transcription factors (TFs) to cognate motifs. In fact, TFs are a major mechanism of dynamic regulatory activity^35,43,44,88^, with activity influenced by epigenetic state of the genomic locus^89,90^ and TF affinity for the binding site^91,92^, among other factors.

We scanned MPRA-emVars (p_FDR_<=0.1) for variant-specific disruptions in predicted TF binding affinity (q_Storey_<0.05) across 505 unique TFs expressed in mature iGLUTs (1,573 motifs) and compared correlations between TF binding disruptions and MPRA variant-specific activity. We annotated significantly correlated TFs based on downstream gene expression changes, cue-specific changes in chromatin accessibility, and differential peak motif-enrichments (**Fig. 4**). Top TF motif enrichments, chromatin accessibility, and TF gene expression varied by context (**Fig. 4A; SI Fig. 16C**) and included expected immune-responsive TFs: IL-6 with interleukin-associated TFs (e.g., CREB1, SMAD3); IFNα with interferon targets (e.g., IRF9, STAT1) (**SI Fig. 16A-B, SI Data 2.20**), as well as immune-responsive TFs involved in neural development and differentiation. For example, binding affinity z-scores of TFAP2C_5, involved in neural development, most highly correlated with activity following exposure to IFNα (r=0.81, p=2×10^-4^, n=18) (**Fig. 4B**).

**Figure 4.**
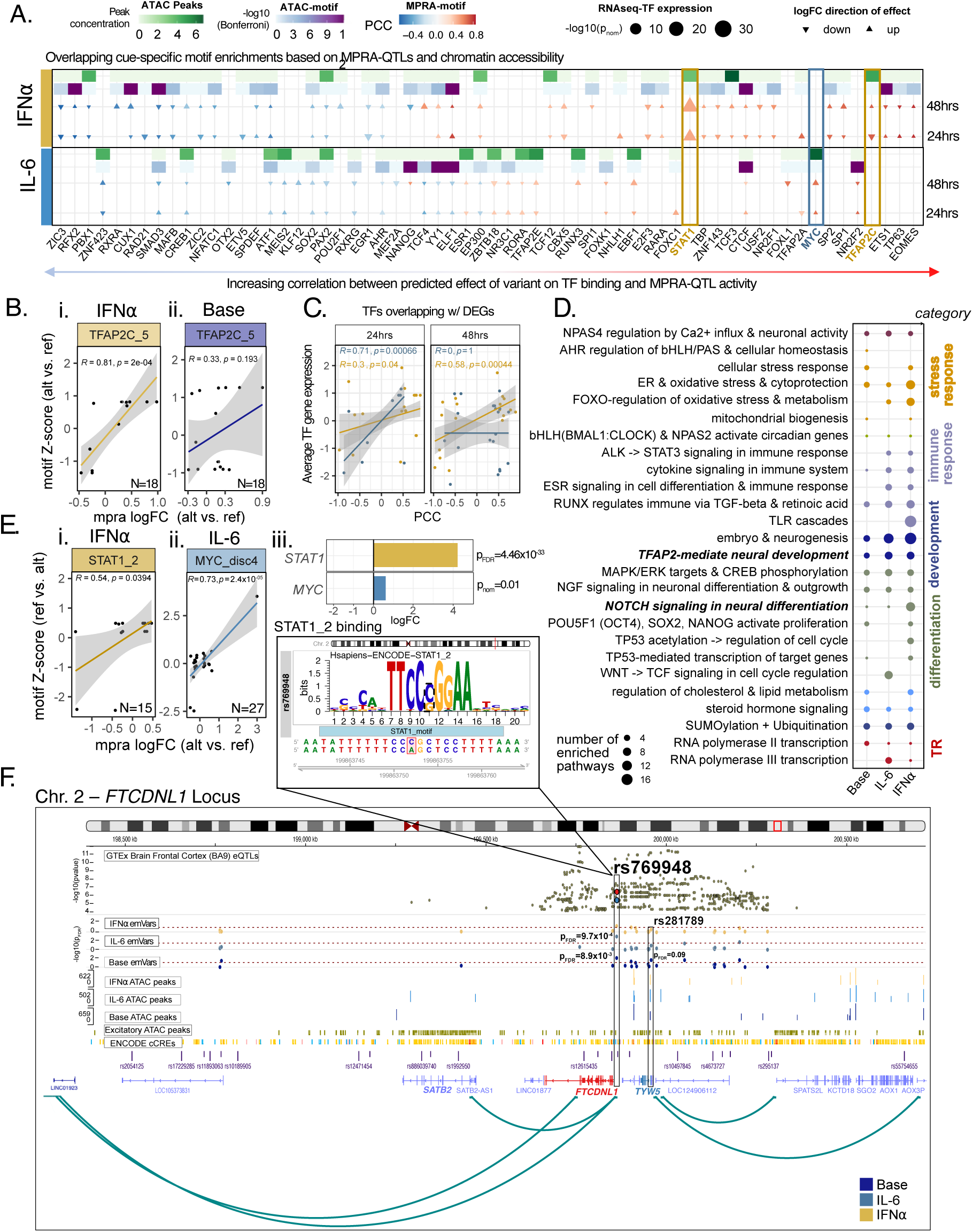
Dynamic regulation associated with variant-specific binding affinities and cue-specific changes in chromatin accessibility, related to SI Fig. 18, SI Data 2.20. Allele-specific effects on transcription factor (TF) binding affinities tested across 1,573 binding motifs representing 505 TFs expressed in mature iGLUTs. TFs regulating MPRA-emVars were identified by integrating measured allelic effects (p_FDR_ <0.1) and affinity binding z-scores (TF motif-SNP interactions (q_Storey_ <0.05). **(A)** Top predicted TFs regulating MPRA-emVars overlapped with cue-responsive ATAC peaks motif enrichments and showed differential cue-responsive gene expression patterns that were temporally dynamic (24hrs-vs-48hrs). Dot plot and heatmap representing top TFs predicted to regulate MPRA-emVars that either overlapped with differentially accessible ATAC peaks (cue-vs-vehicle) (green; color=peak concentration), ATAC motif enrichments (purple; color=-log10(p_Bon_)) or differentially expressed genes (size=-log10(P-value), shape=direction of logFC (up or down regulated), with the Pearson’s Correlation Coefficient as the fill of each dot (blue=negative correlation; red=positive) and TFs organized by magnitude of correlation. **(B)** TFAP2C_5 is a top correlated motif following IFNα exposure (r=0.81, p=2×10^-04^, n=18), but not at baseline (r=0.33, p=0.193, n=18); individual SNPs represented as points (R=Pearson’s Correlation Coefficient). (C) TF average gene expression was significantly positively correlated with MPRA-motif correlations at 24hrs (IL-6: r=0.71, p=0.00066, IFNα: r=0.3, p=0.04) and 48hrs (IFNα: r=0.58, p=0.00044)**. (D)** Comparative over-representation analysis (REACTOME pathways) between baseline, IL-6, and IFNα conditions (TFs selected from **A**). Size represents the total number of unique term enrichments affecting a signaling cascade, ordered and colored by broader category. Signaling cascades related to example TFs (B, E) are in bold. **(E)** Known downstream TFs of interferon and interleukin signaling regulate MPRA-emVars. (**i**) STAT1_2 (Pearson’s r=0.54, p=0.0394, IFNa emVars n=15, interaction emVars n=2) and (**ii**) MYC_disc4 (Pearson’s r=0.73, p=2.4×10^-5^, n=27, interaction emVars n=4) were highly correlated with MPRA variant-specific effects, and (**iii**) *STAT1* (logFC=4.3, p_FDR_=4.46×10^-33^) and *MYC* (logFC=0.58, p_nom_=0.01) were differentially expressed following IFNa and IL-6 exposure respectively. **(F)** The *FTCDNL1* locus, associated with cognition, SCZ, and metabolic syndrome by GWAS, is dynamically regulated by immune responsive MPRA-emVars (rs769948, rs281789) with predicted proximal and distal gene targets. The reference allele of both rs769948 (C) and rs281789 (T) were predicted to increase affinity of STAT1 binding (rs769948 q_Storey_= 0.00085; rs281789 q_Storey_= 0.037). The rs769948 reference allele (C) significantly increased transcriptional activity at baseline and with IL-6 (mpralm; Base: p_FDR_=8.9×10^-3^; IL-6: p_FDR_=9.7×10^-4^) but was repressed following IFNα exposure. rs281789 was a significant MPRA-emVar only at baseline (base p_FDR_=0.09, IL-6-v-base p_FDR_=0.028, IFNa-v-base p_FDR_=0.037). Both overlap with annotated brain regulatory elements (ENCODE) and iGLUT-specific chromatin loops (track) and are brain-eQTLs (GTEx v8 Frontal Cortex) of *FTCDNL1* and *TYW4*. Integration of iGLUT specific chromatin accessibility peaks and Hi-C loops (ABC) further map these dynamic emVars to regulation of distal targets >500KB away, including *STAB2* (rs769948) and *AOX1*/*AOX3P* (rs281789).

MPRA variant effects were positively correlated with cue-responsive TFs expressed in iGLUTs (strongest at 24hrs post exposure for IL-6; R=0.71, p=0.00066, 48hrs post exposure for IFNα; R=0.58, p=0.00044) (**Fig. 4C**). Comparative gene set enrichment analysis of the TFs implicated in regulating responses at baseline and in response to IL-6 and IFNα resolved interleukin, STAT, and immune signaling as expected, but also highlighted enrichments for cellular stress response, metabolism, and neural differentiation (**Fig. 4D**).

For example, predicted disruptions in motif binding affinities of STAT1 (r=0.54, p=0.0394, n=15) and MYC (r=0.73, p=2.4×10^-5^, n=27) , both involved in NOTCH signaling cascades impacting neural differentiation, were highly correlated with context-responsive MPRA-emVars effects (**Fig. 4E,i-ii**), differentially enriched in context-specific ATAC accessible regions (STAT1, IFNα*: q_Storey_=0.08,* MYC, IL-6*: q_Storey_=0.02),* and corresponded to dysregulated TF expression (STAT1, logFC=4.5, p_FDR_=4.46×10^-33^; MYC: logFC=0.58, p_nom_=0.01; **Fig 4Eiii**).

Dynamic transcriptional regulation (MPRA-emVars), chromatin accessibility (ATAC peaks), chromatin contact (iGLUT Hi-C loops), and variant-specific predicted TF binding affinities influenced dynamic genotype-by-immune interactions at psychiatric GWAS loci (**Fig. 4F**). For example, the *FTCDNL1* locus, associated with SCZ, metabolic syndrome, and cognition, was dynamically regulated by MPRA-validated brain-eQTLs with proximal and distal gene targets: the reference alleles at *rs769948* (C) and rs281789 (G) were predicted to increase binding affinity for STAT1 (q_Storey_=0.0008, q_Storey_=0.037, **Fig. 4F**). *rs769948*-C significantly increased transcriptional activity at baseline (p_FDR_=8.9×10^-3^) and with IL-6 (p_FDR_=9.7×10^-4^), but was repressed following exposure to IFNα, whereas *rs281789-G* increased activity only at baseline (base: p_FDR_=0.09, interaction IFNα-v-base p_FDR_=0.057, IL6-v-base p_FDR_=0.03). While brain eQTLs mapped these SNPs to regulation of *FTCDNL1/TYW4*, enhancer-to-gene mapping with iGLUT-specific chromatin loops also predicted multi-gene regulation at *STAB2* (∼500KB upstream; schizoaffective disorder and serum level immunoglobulin glycosylation GWAS target gene) and *AOX1/AOX3P* (∼750 kb downstream; associated with response to antidepressants and drug metabolism) (**Fig. 4F**). Overall, dynamic regulation of GWAS risk loci in response to neuronal inflammation is associated with variant-specific binding affinities and cue-specific changes in TF gene expression and chromatin accessibility.

## Discussion

By testing the regulatory activity of 152 GWAS loci linked to brain diseases and traits across dynamic immune-regulated cues in live human neurons, we modelled neuronal immune response during neurodevelopment, demonstrating gene x environment interactions of broad relevance to subsequent neuropsychiatric outcomes. We characterized effects at 152 GWAS loci, testing 3,668 CRSs in human neurons, of which 1,156 were active in at least one condition. In total, we resolved 267 variant-specific MPRA-emVars across 859 (baseline), 860 (IL-6), and 719 (IFNα) active MPRA-CRS; 31% had significant cue-by-variant interaction effects (interaction MPRA-emVars). Overall, dynamic effects tended to affect the magnitude but not the direction of regulatory activity. Our neuron lenti-MPRA replicated MPRA-CRS previously reported in NPCs, was concordant with brain eQTLs, and highlighted known risk-associated eGenes, but expanded upon these existing resources by incorporating dynamic regulatory activity.

Fetal brain development is exquisitely sensitive to inflammatory insults throughout pregnancy^93^, with both maternal^17^ and fetal^93^ immune dysfunction, as well as genetic defects resulting in type I IFN overproduction^94,95^, linked to altered neurodevelopment and increased risk of psychiatric and neurological outcomes in offspring. We selected two cytokines, IL-6 and IFNα, based on evidence from animal models^20,21,96,97^, cell culture^28-^_30,59_, inflammatory biomarkers^98–100^, brain imaging^101,102^, and post-mortem brain analyses^26,103^. Both cytokines are believed to act directly on neurons: maternal IL-6 accumulates in the placenta^104^ and crosses the fetal blood brain barrier^105^, whereas IFNα is generated by fetal brain cells in response to infection^31^.

Despite being a simple acute treatment paradigm, we observed time-dependent effects worth considering in subsequent studies of dynamic genetic regulation. First, *IL6R* and *IL6ST* expression increased within 24-hours of IL-6 treatment but declined nearly to baseline by 48-hours; likewise, *IFNΑR1* and *IFNΑR2* expression decreased following exposure to IFNα, but returned to baseline by 48-hours. These trends paralleled genome-wide effects; although gene expression signatures were strongly correlated across time points, there were fewer DEGs and decreased magnitude of effects over time, consistent with a rapid initial response and subsequent attempt to return to homeostasis (of note, mature iGLUTs were treated one time, without re-exposure). Transcription at 24-hours post-treatment correlated more strongly with MPRA effects measured at 48-hours. These dynamic patterns indicated that gene expression and chromatin accessibility changes should be monitored over time, with careful consideration of pseudotime effects in MPRA studies when dissecting mechanistic pathways impacted by cue-dependent genetic regulation.

Our library design prioritized eQTLs from the adult brain, but MPRA was conducted in iGLUTs that more resemble fetal-like neurons; the resulting MPRA-emVars showed significant concordance with both adult and fetal eQTLs. Patterns of gene expression change across neurodevelopment^106,107^ and aging^108^, which may underly critical periods of heighted susceptibility to environmental stressors. Such changes presumably arise from differences in genetic regulation, reflecting shifting patterns of TF expression that mediate enhancer activity^109,110^. Age-specific eQTLs modify the functional impact of genetic risk throughout lifespan^37,111,112^ and are particularly relevant given the variable age of onset of symptom presentation across brain disorders. A variety of methods now exist to maintain^113,114^ or accelerate^115–118^ aging *in vitro*, which could be incorporated into future MPRAs, towards dissecting the dynamic functional impact of genetic risk across neurodevelopment, brain maturation, and aging to inform mapping of brain-related GWAS and facilitate precision medicine.

Notable technical limitations in MPRA design and methodology reduce the broad generalizability of our GxE analyses. First, reflecting the present state of GWAS, the variants tested were identified in exclusively European-ancestry data and excluded the MHC locus. Moreover, many recent publicly available GWAS remain underpowered, overall biasing the MPRA library towards specific disorders (e.g., eating disorders, PTSD and OCD had very few variants included). Second, technical limitations restricted CRSs to short DNA fragments flanking prioritized variants, potentially omitting crucial portions of larger regulatory regions. Recent technological innovations in MPRA (e.g., tiling MPRA design^119^) should facilitate examination of more complex regulatory sequences moving forward. Third, although we designed the library to test 8,960 sequences, of which 7,267 were represented with adequate barcode coverage in our physical lenti-MPRA library. Of these, we acquired reads for ∼6,700, filtered to 3,668 with adequate barcode coverage.

MPRA design should be mindful of physical library complexity and neuronal culture sizes to ensure comprehensive analysis of their full MPRA library. A notable limitation of lenti-MPRA design is the risk of CRS-barcode swapping, mitigated here by our use of 5’UTR barcoding. Fourth, although MPRA activity was highly correlated between donors (r=0.87-0.95), polygenic donor genotype may influence functional genomics, especially when exploring different contexts, and warrants further exploration^120,121^. Fifth, while MPRA-emVar effects were correlated with previous findings, the number of overlapping sequences was small (<10%) and do not provide robust validation; four previously published MPRAs^54,78,122,123^ based on psychiatric GWAS likewise had low to moderate overlap in variants (**SI Data 2.17**). Lastly, as CRSs were tested outside their endogenous context, results may not accurately inform regulatory activity of distal gene targets. Although new methods to integrate matched cue-specific epigenetic datasets refine predictions of gene targets of enhancer activity^85,124^, further experimental validation through crisprQTL^125^ or prime-editing^126^ is warranted. Future studies of cue-specific GxE effects across additional variants, doses, longitudinal and recovery time-points, cues, cell-types (particularly brain-specific immune cells), and ultimately within more physiologically relevant brain organoids via emerging single-cell MPRA methods^127^, and across an expanded number of donors via village-in-a-dish approaches^128^ will be crucial for further dissection of how immune activation during fetal development contributes to brain disorder risk.

The influence of pro-inflammatory environmental factors on brain-related regulatory elements may explain biological mechanisms through which immune signaling contributes to increased susceptibility for complex brain disorders. We identified hundreds of GWAS variants that confer greater susceptibility to complex brain disorders following developmental exposure to neuroinflammation and mapped cue specific risk-associated genes, informing the influence of GxE interactions across complex brain traits. Dynamic neuronal immune-response genetic regulation frequently reflected combinatorial effects of multiple variants within a GWAS loci, with distinct variants top ranked across neuroimmune cues; moreover, loci with immune-specific genetically regulated transcription were annotated for pleiotropic effects across multiple traits. Likewise, downstream target genes of interaction MPRA-emVars were uniquely enriched for complex brain disorders and common comorbid metabolic and immune syndromes. The clinical impact of resolving GxE interactions include preventative measures (e.g., improved prenatal care, social supports, and early life interventions for high-risk individuals) and care of current patients (e.g., drug repurposing, patient stratification by immune status, personalized prescription). Overall, when attempting to understand the genetic mechanisms of variable penetrance and pleiotropy, with broad relevance across complex traits and disorders, it is critical to consider the impact of dynamic regulation of gene expression.

## Methods

### Lenti-MPRA library design across neuropsychiatric trait GWAS

A Lenti-MPRA^129^ library was designed by statistical fine-mapping of ten GWAS: (Alzheimer’s disease (AD)^67^, attention deficit hyper-activity disorder (ADHD)^68^, anorexia nervosa (AN)^69^, autism spectrum disorder (ASD)^3^, bipolar disorder (BIP)^70^, major depressive disorder (MDD)^71^, obsessive compulsive disorder (OCD)^72^, post-traumatic stress disorder (PTSD)^73^, schizophrenia (SCZ)^1^, and neuroticism (NEU-P)^74^) using two complimentary methods incorporating dorsolateral prefrontal cortex (DLPFC)^75^ expression quantitative trait loci (eQTLs): Bayesian co-localization (coloc2^76^) and transcriptomic imputation (S-PrediXcan^77^) (**SI Fig. 7A; SI Data 2.1-2.3**). For the former, significant GWAS loci and Dorsolateral Pre-frontal Cortex (DLPFC)^75^ eQTLs from the Common Mind Consortium (CMC) were tested for co-localization using a lenient significance threshold for GWAS loci (p<1×10^-6^). The most probable causal eQTLs from moderately to highly colocalized loci (PPH4>=0.5) were selected, along with all SNPs in high LD (r^2^ >= 0.9) (**SI Data 2.2-2.3**); controls for coloc2 prioritization were selected from significant BIP GWAS loci that (i) did not colocalize (PPH3 > 0.9) and (ii) were not significant CMC DLPFC eQTLs. For S-PrediXcan, all SNPs within the predictor models of Bonferroni-corrected significant (p<4.64×10^-6^;(0.05/10786)) trait-associated S-PrediXcan genes (eGenes) and all SNPs in high LD (r^2^ >= 0.8) with them were selected (**SI Data 2.4-2.5**).

Several strategies were applied to incorporate appropriate controls. First, scramble sequences (n=100) were added as negative controls. Only the scramble controls were used for normalization and as a measure of basal activity of the minimal promoter. For a point of comparison, two additional sets of benchmark variants were included. As empirical controls, 310 SCZ and AD GWAS SNPs producing the greatest (n=164) and least (n=146) transcriptional shifts in K562 chronic myelogenous leukemia lymphoblasts and SK-SY5Y human neuroblastoma cells^78^ that overlapped with CMC DLPFC eQTLs were included to compare results of previously tested sequences. Third, to query the regulatory activity of significant GWAS loci lacking eQTL associations, we included 88 significant BIP GWAS SNPs that (i) did not colocalize (PPH4<0.1/PPH3>0.9) and (ii) were not significant CMC DLPFC eQTLs.

After accounting for SNPs identified through multiple strategies, and removal of SNPs with sequences containing restriction digest sites (AgeI/SfbI), we synthesized a library of 4,430 SNPs (8,860 variants), many of which overlapped with chromatin accessible peaks in the DLPFC and hiPSC-derived neurons, and 100 scramble controls. Oligonucleotides were synthesized by Agilent and cloned into the lenti-MPRA vector ^52^.

We performed a power analysis using designmpra^130^, which estimates the power of a t-test to distinguish differential activity in a variant at a Bonferroni corrected alpha=.05 level. At 4430 bi-allelic pairs, 100 barcodes per SNP, activity standard deviation of 1 (typical range = 0.3-2), and 4 replicates, the power of a t-test to detect differential variant shifts of (logFC) of 0.5 or greater at a Bonferroni corrected alpha=0.05 level is 100%.

### Lenti-MPRA library preparation and viral titration

The MPRA library was generated according to published lenti-MPRA protocols with slight modifications^129^. Briefly, 200 base pair oligonucleotides flanking each prioritized SNP were synthesized by Agilent to create an MPRA library of 9,244 neuropsychiatric associated variants. The Agilent oligo pool was PCR amplified, and a minimal promoter and spacer sequence added downstream of the CRS. This protocol uses a 5′ UTR barcoding method uses a shorter distance (102 bp) between the CRS and barcode than 3′ UTR barcoding methods (801 bp), reducing the risk of CRS–barcode swapping. Amplified fragments were purified and amplified again for 15 cycles to add a random 15bp sequence to serve as a unique barcode. Barcoded fragments were inserted in the *Sbf*I/*Age*I site of the pLS-SceI vector (AddGene #13772) and then transformed into 10-beta competent cells (NEB, C3020) via electroporation. Bacterial colonies were grown overnight on Ampicillin-positive plates and midi-prepped for plasmid collection. The quality of the purified plasmid was evaluated by Sanger Sequencing of 16 colonies at random. CRS-barcode associations were identified by sequencing of the purified plasmid (MiSeq; paired-end; 15milion reads). Our final MPRA library consisted of 7,267 CRS (77% of the designed) that were represented at a minimum of 10 barcodes. 2^nd^-generation lentiviral packaging of the purified plasmid was performed by the viral core at Boston’s Children Hospital. To determine MOI and approximation of appropriate viral volume we infected day 14 iGLUTs (0, 1, 2, 4, 8, 10, 16, 32, 64 µL) with control lentivirus (pLS-SV40-mP-EGFP; AddGene #137724) and harvested for 48hrs later. Following DNA isolation, we performed qPCR to calculate the MOI based the relative ratios of genomic DNA to inserted viral DNA (after subtracting background noise caused by residual backbone DNA).

### Cell-type deconvolution of postmortem DLPFC

We checked for the relative abundance of cell-type in the postmortem CMC DLPFC by performing cell-type deconvolution with the R package dtangle^131^ and a single-cell expression reference panel from the cerebral cortex. In the CMC DLPFC, glutamatergic neurons make up the highest proportion of cells based on cell-type deconvolution.

### NGN2-glutamatergic neuron induction from clonalized hiPSC lines for molecular experiments^56,57^

Clonal hiPSCs from two neurotypical donors of European ancestry with average schizophrenia PRS and no history of psychiatric diagnoses (#3182 (XX) and #2607 (XY)) were generated by lentiviral transduction with pLV-TetO-hNGN2-eGFP-Neo and lentiviral FUW-M2rtTA (Addgene #20342), followed by antibiotic selection and clonal expansion. Stably selected clones were validated to ensure robust cell survival, expression of fluorescent tags, and transgene expression. hiPSCs were maintained in StemFlex™ Medium (ThermoFisher #A3349401) and passaged with EDTA (Life Technologies #15575-020).

On day 1, medium was switched to non-viral induction medium (DMEM/F12 (Thermofisher, #10565018), 1% N-2 (Thermofisher, #17502048), 2% B-27-RA (Thermofisher, #12587010)) and doxycycline (dox) was added to each well at a final concentration of 1 µg/mL. At day 2, transduced hiPSCs were treated with 500 µg/mL G418 (Thermofisher, #10131035). At day 4, medium was replaced including 1 µg/mL dox and 4 µM cytosine arabinoside (Ara-C) to reduce the proliferation of non-neuronal cells. On day 5, young neurons were dissociated with Accutase Cell Detachment Solution (Innovative Cell Technologies, # AT-104), counted and seeded at a density of 1×10^6^ per well of a Matrigel-coated 12-well plate. Medium was switched to Brainphys neuron medium (Brainphys (STEMCELL, # 05790), 1% N-2, 2% B27-RA, 1 μg/mL Natural Mouse Laminin (Thermofisher, # 23017015), 10 ng/mL BDNF (R&D, #248), 10 ng/mL GDNF (R&D, #212), 500 μg/mL Dibutyryl cyclic-AMP (Sigma, #D0627), 200 nM L-ascorbic acid (Sigma, # A4403)). For seeding, 10 mM Thiazovivin (Millipore, #S1459), 500 μg/mL G418 and 4 μM Ara-C and 1 μg/mL dox were added. At day 6, medium was replaced with Brainphys neuron medium with 4 μM Ara-C and 1 μg/mL dox. Subsequently, 50% of the medium was replaced with fresh neuronal medium (lacking dox and Ara-C) once every other day until the neurons were harvested at D24.

### *Cue-Specific RNAseq* (SI Fig. 1, 2-5, 8; SI Data 1)

Briefly, mature iGLUTs (22 DIV), were acutely exposed (24 or 48 hours) to IL-6 (25 ng/mL (24hr only) and 60ng/mL), IFNα (100 UI/mL (24hr only) and 500 IU/mL), or vehicle (0.1% FBS in ultrapure H_2_O before harvest at 24 DIV (two control donors, three to four replicates per donor per cue/vehicle treatment; experimental schematic: **SI Fig. 1**).

| Compound | Solvent | Supplier | Product # | Conc. |
| --- | --- | --- | --- | --- |
| Human-recombinant IL-6 | UltraPure H<sub>2</sub>O | Sigma/Aldrich | GF338 | 60 ng/mL |
| Human-recombinant INFα2-b | UltraPure H<sub>2</sub>O | Mount Sinai Pharmacy | NDC 0085-4350-01 | 500 IU/mL |

RNA Sequencing libraries were prepared using the Kapa Total RNA library prep kit. Paired-end sequencing reads (100bp) were generated on a NovaSeq platform. Raw reads were aligned to hg38 using STAR aligner^132^ (v2.5.2a) and gene-level expression were quantified by featureCounts^133^ (v1.6.3) based on Ensemble GRCh38 annotation model. Genes with over 10 counts per million (CPM) in at least four samples were retained. Surrogate variable analysis (SVA) was performed to assess the correlation of known covariates and identify surrogate variables contributing to variance using the R packages sva^134^ and variancePartition^135^ (e.g. batch, treatment, donor, replicate, sex, and RIN). Following identification of SVs and covariates to correct for in the model, raw read counts were normalized with voom^102^ and, due to the repeated measures study design, where individuals are represented by multiple independent technical replicates, differential expression analysis with repeated measures was performed by the *Dream* method from variancePartition^135^. Bayes shrinkage (limma::eBayes) estimated modified t- and p- values. Gene level significance values were adjusted for multiple testing using the Benjamini-Hochberg method to control the false discovery rate (FDR). Genes with FDR < 5% were considered significantly differentially expressed (limma::TopTable)^136^ (**SI Data 1.1-1.5**). In these analyses, the t-test statistics from the differential expression contrast were used to rank genes in the GSEA using the R package ClusterProfiler^106^ (**SI Data 1.6**). Gene-set enrichment analysis using WebGestalt^137^ was performed between p_nom_<0.05 DEGs and gene-sets curated from four rodent models of maternal immune activate: poly(I|C) exposure (conceptus, whole brain, amygdala, and frontal cortex)^24,66,138^, IL-6 exposure (whole brain)^24^, *H1N1* Flu virus infection (whole brain)^24^, and chronic unpredictable maternal stress (whole brain)^139^.

### Meta-analysis of gene expression across cues

We performed a meta-analysis and Cochran’s heterogeneity Q-test (METAL^140^) using the p-values and direction of effects (t-statistic), weighted according to sample size across all sets of perturbations (Target vs. Scramble DEGs). Genes were defined as convergent if they (1) had the same direction of effect across cue exposure (2) were Bonferroni significant in our meta-analysis (Bonferroni adjusted p-value; p_Bon_<= 0.05), and (3) had a non-significant Cochran’s Heterogeneity Test (**SI Data 1.7**).

### *Cue-Specific ATAC-Seq* (SI Fig. 1, 4-5; SI Data 1)

Briefly, mature iGLUTs (22 DIV), were acutely exposed (48 hours) to IL-6 (60ng/mL), IFNα (500 IU/mL), or vehicle (0.1% FBS in ultrapure H_2_O)before harvest at 24 DIV (two control donors, two replicates per donor per cue/vehicle treatment; experimental schematic: **SI Fig. 1**). Mature neurons were washed with 500uL of PBS (-Ca/-Mg)-0.5mM EDTA per well of a 12-well plate. Then, 300uL dissociation solution (0.042 U/µL papain suspension (Worthington-Biochem LS003126) in HBSS (Thermofisher #14025076)-10mM HEPES (Thermofisher #J61275AE)-0.5mM EDTA (Life Technologies #15575-020), pre-activated at 37C for 5 minutes) supplemented with 0.017U/µL DNase (Thermofisher #EN0521) and 1x Chroman I was added to each well before incubating the plate at 37C for 10 minutes, shaking at 125rpm. 600uL deactivating solution (DMEM-FBS-Chroman I) was then added to each well, and cells were dissociated into single cells by pipetting gently. For each condition, cells from 4 wells of a 12-well-plate were combined into a single 15mL conical tube for higher yield. After spinning at 600g for 5 minutes at room temperature, cells were resuspended in 310uL of DMEM (Thermofisher #10566-016)-10% FBS. Then, the cell suspension was filtered through a 37um reversible strainer and frozen in DMEM-10% FBS-10% DMSO.

ATAC sequencing library prep and sequencing were performed by the Yale Sequencing Core. The adaptor sequence for pair-end sequencing was removed using trim_galore^141^ and sequencing quality measure by FastQC^142^ and MultiQC^143^. Each experiment contained two technical replicates and two biological replicates (four samples total). All R1 and R2 fastqs passed basic QC with FastQC; total sequences per sample ranged from 47.2 million to 79.7 million after removal of mitochondrial reads (average 61M), with percent deduplicated ranging from 65%-80% (average 74.38%). Average GC content was normally distributed with a mean of 44% and average sequence length 85-119. Data was aligned with Bowtie2^144^ against hg38 reference genome including rare SNVs. Mitochondrial reads were removed, sam files sorted and indexed, and converted to compressed BAM files with samtools^145^. Peak calling and reproducibility analysis were performed using MACS2 (2.1.0)^146^ and IDR (version, soft-idr-threshold 0.05)^147^ respectively, according to ENCODE ATAC-seq pipeline (v1 2019)^148^ specifications. Fraction of reads in peaks (FRiP) scores were calculated for each sample from the alignment file (bam) and MACS2 narrowPeak files: FRiP scores ranged from 0.23-0.35, passing ENCODE ATACseq data standards (**SI Data 1.8**). All replicates passed the two ENCODE standards for IDR, with ratio of pooled pseudoreplicate results to true replicate results (Np/Nt) as well as self-replicate peaks (N1/N2) within a factor of two. Self-consistency ratio: max(N1,N2)/ min(N1,N2) = 1.56-2.86; and Rescue Ratio: max(Np,Nt)/ min(NP,Nt)=1.32-1.64.Consensus peak sets were defined as ATAC peaks present in at least 4 out of 6 pairwise true-replicate IDR comparisons for each condition (4 samples total), with at least 50% overlap (bedtools intersect -wa -u -f 0.50) resulting in a range of 19,373-27,930 peaks per condition. A single merged file was generated for each condition for visualization purposes by merging bam files with samtools, then intersecting that file with the list of consensus peaks via bedtools intersect. Bed files for each merged peak were opened with Integrative Genomics Viewer (IGV_2.16.0)^149^ for figure generation. Separate narrowpeak files for each ATAC replicate were also intersected with the list of consensus peaks per condition to be used for differential accessibility analysis with DiffBind^150^. Counts for all replicates across conditions were normalized to full library size, donor was used as a blocking factor, treating within donor replicates as repeated measures, condition-vehicle comparisons were defined as individual contrasts, and differential sites for each contrast calculated using edgeR at a threshold of Benjamin-Hochberg method of multiple testing correction; p_FDR_<0.1, or a nominal p-value <0.05 for subsequent analysis. Transcription factor binding site motif enrichment was performed using Homer (findMotifsGenome.pl -size given -mask) on differential accessible regions identified at both significance thresholds^151^. Peak annotation was performed using ChIPseeker^152^ (v 1.8.6) and gene set over representation analysis of genes mapped to open chromatin regions was performed with ClusterProfiler^153^ using Gene Ontology, WikiPathway, KEGG, and REACTOME gene-sets.

### *Cue-specific MPRA* (SI **Fig. 1**, 7-18; SI Data 2)

Briefly, the MPRA library was transduced into mature iGLUTs (21 DIV), 24 hours later neurons were acutely exposed (48 hours) to IL-6 (60ng/mL), IFNα (500 IU/mL) or vehicle (0.1% FBS in ultrapure H_2_O) before harvest at 24 DIV (two control donors, two biological replicates each, experimental schematic: **SI Fig. 1**). Day 21 iGLUTs were spinfected (1krcf for 1 hr @37C, slow accel, slow deceleration) with lenti-MPRA library, (based on titrations of the control virus from the Gordon et al. 2020 lenti-MPRA Nature Protocol Supplement). 24 hours after spinfections, full media was replaced to remove un-integrated virus. At 48 hours post-infection, iGLUTs were treated with stress and inflammatory cues or basal media. The number of cells required pre-replicate was calculated according to the Gordon et al. 2020 protocol, with on average, 2 million hiPSCs seeded per well of a six-well plate and later batched together as one technical replicate for an average of 6 million cells per replicate. 72hrs post-lentiviral-infection, and 48 hours post-exposure to stress or inflammatory compounds neuronal cells were washed three times and harvested using AllPrep DNA/RNA mini kit (Qiagen) and the libraries prepped as previously described. The libraries were sequenced as paired end reads on a NextSeq 2×50 on S2 flow cell (3.3-4.1 B reads/cell) by the New York Genome Center.

### Lenti-MPRA CRS-barcode association from MiSeq

Sequencing of purified plasmid DNA was sequenced on an Illumina MiSeq v.2 (15 million paired-end reads and generated fastq files with bcl2fastq (parameters: --minimum-trimmed-read-length 0 --mask-short-adapter-reads 0). Barcode-CRS association was performed as previously described using the association utility of MPRAflow v2.3.5^129^ (run as: nextflow run association.nf – fastq-insert –fastq-bc –fastq-insertPE –mapq 3 –baseq 15).

### Lenti-MPRA RNA/DNA counts

We demultiplexed the indexed DNA and RNA libraries and generated paired-end fastq files with bcl2fastq v2.20 and used the count utility of MPRAflow 2.3.5 both with and without the --mpranalyze flag included (run as: nextflow run count.nf -w –experiment-file –dir –outdir –labels –design –bc-length 14 –umi-length 16 --thresh 10) to compute the activity score for each element and produce count files formatted for analysis with MPRAnalyze (**SI Data 2.10**)^79^. Roughly 6,700 inserts were sequenced (92% of the library). We filtered out barcode-CRS pairs with low DNA counts (<15) and those with RNA counts, but no DNA counts, leaving 3,668 CRSs with a minimum requirement of 10 barcodes each.

### Lenti-MPRA comparison of regulatory activity across replicates, donors, and cue exposures

Transcriptional activity, measured as the normalized log2 DNA and RNA counts per CRS, across conditions was strongly correlated between replicates (Pearson’s Correlation Coefficient r=0.98-1.00). Normalized log2(RNA/DNA) ratios were also strongly correlated (mean r=0.83, max r=0.93), with only 1 comparison (donor 2607 following IFNα exposure below a correlation of 0.83; at 0.61; **SI Fig. 8)**. When technical replicates were averaged together and compared across donors, correlations were high (r=0.87-0.95) and we retained all replicates in downstream analyses. (**Fig. 1A; SI Data 2.11**). Across sequences the number of barcodes per unique CRS were highly correlated between CRS shared across all conditions (r=0.997-0.999). There was no significant difference in mean number of barcodes (n∼45) per insert or the proportion of CRS by prioritization method or disorder association across the conditions.

After filtering we re-performed a power analysis (designmpra^130^) with the actual values post filtering. At 1000 bi-allelic pairs, with an average of 45 barcodes per SNP, activity standard deviation of 1 (typical range = 0.3-2), and 4 replicates, we have 80% power to detect differential variant shifts of (logFC) of ∼0.5 at a Bonferroni corrected alpha=0.05 level using a t-test.

### Comparison of three approaches for evaluating MPRA activity and variant specific shifts

We performed comparison of quantification, variant specific differential analysis, and cue-by-variant interaction testing across three methods: MPRAnalyze^79^, mpralm^80^, and DEseq2^81,82^:

### MPRAnalyze^79^

#### Quantification

Mean transcriptional activity across replicates was calculated using MPRAnalyze. First, library size correction factors were estimated across replicates and donors using estimateDepthFactors(obj, lib.factor=c(“replicate”,”donor”,) which.lib=”both”, depth.estimator=”upper quantile”). Activity was quantified using analyzeQuantification (obj = obj, dnaDesign = ∼ donor + replicate, rnaDesign = ∼ donor + replicate). Alpha values representing the transcriptional rate of each sequence were extracted from the fitted model. CRS were labeled active if the MAD score p_FDR_<=0.05.

#### variant-specific differential analysis

First, RNA/DNA counts were separated by reference and alternative alleles and merged. Library size correction factors were estimated across replicates and donors using estimateDepthFactors(obj, lib.factor=c(“replicate”,”donor”) which.lib=”both”, depth.estimator=”upper quantile”). Differential activity was calculated between alternate (alt) and reference (ref) alleles using the classic mode version of analyzeComparative(mpraobject, dnaDesign = ∼ replicate + donor + variant + barcode, rnaDesign= ∼ variant, reducedDesign= ∼ 1, mode=”classic”) and testLrt(obj).

#### cue-by-variant interaction analysis

For active MPRA CRS and significant MPRA-emVars at either baseline or in IL-6 or IFNα, cue-by-variant effects were tested using an interaction analysis. First, conditions were merged based on shared sequences and library size correction factors were estimated across replicates, donors, and conditions using estimateDepthFactors(obj, lib.factor=c(“replicate”,”donor”,”cue”) which.lib=”both”, depth.estimator=”upper quantile”). Possible interaction effects between IL-6 or IFNα exposure and baseline (vehicle) was performed using a cue-by-variant interaction term and significance tested using analyzeComparative(mpraobject, dnaDesign= ∼ replicate + donor + cue_variant, rnaDesign= ∼ cue_variant, reducedDesign= ∼ 1, mode=”classic”) and MPRAnalyze::testLrt(obj)”.

### mpralm^80^

#### quantification

We assessed the activity of CRS compared to the average activity of all CRS with *mpralm()* (mpra v.1.16.0) followed by *eBayes() abd topTable()* (limma v3.50.3) after correcting for donor and replicate (∼ 1 + donor + replicate). Sequences that had positive logFC (>0) and a met a statistical significance threshold at FDR < 0.1 were labeled as “active CRS” while those with negative logFC and FDR<0.1 were labeled as “repressed CRS”. Only “active CRS” were used in downstream analysis of differential variant effects.

#### variant-specific differential analysis

We applied *mpralm()* (mpra v.1.16.0) followed by *topTable()* (limma v3.50.3) to detect allelic effects in MPRA active CRS (MPRA sequence pairs with one of sequence active in any condition). We set a statistical significance threshold at FDR < 0.1 to define emVars.

#### cue-by-variant interaction analysis

Cue-by-variant interaction effects were assessed with ∼ cue:variant + donor + rep. Only MPRA CRS with significant allelic differences (MPRA-emVars) in at least one condition were used to test for interaction effects. We set a statistical significance threshold at FDR < 0.1 to define interaction emVars.

#### DEseq2

CRS quantification and allelic analysis with DEseq2 as described^82^:

#### Quantification

We assessed the activity of CRS compared to the average activity of all CRS with DESeq2 using a nested model: ∼ material + material:donor + material:replicate. Here, ’material’ describes the aggregate allele-independent effect observed in RNA compared to DNA (“expression effects”). ’material:donor’ and ’material:replicate’ describes the sample-specific effects nested within RNA/DNA. Active CRS were defined as those with logFC>0 and FDR<0.1.

#### variant-specific differential analysis

*For sequence pairs where at least one pair was an active CRS, we tested for variant specific effects with DEseq2 using a nested model:* ∼ material + material:donor + material:replicate + material:allele. Here, ’material:allele’ describes the allele-specific effects nested with RNA/DNA. We set a 10% FDR threshold to define MPRA-emVars.

#### cue-by-variant interaction analysis

For emVars that were significant in at least one condition, we tested for cue-by-variant interaction effects with DEseq2: ∼ material + material:donor + material:replicate + material:allele + material:allele:cue. Here, ’material:allele:cue’ describes the cue-by-variant interaction effects nested with RNA/DNA. We set a 10% FDR threshold to define interaction emVars.

We compared quantification results across methods and observed that 43% of “active” CRS were identified by all three methods, with the greatest overlap (∼93%) between mpralm and DEseq2. MPRanalyze identified 463 CRS that were not indicated as being active by either mpralm or DEseq2. We also compared the logFC and -log10(p-values) for each analysis across methods (**SI Figure 9-10**). DEseq2 and mpralm methods had the most agreement, with Pearson correlations between 0.9-0.97 for log fold change estimates and correlations between 0.8-0.91. for the -log10(p-value). MPRAnalyze showed the least agreement, moderately correlated with both mpralm and DEseq2 (logFC R=0.59-0.67, -log10(p-value) R=0.48-0.68). We calculated expected p-values for each method and compared them with the observed p-values to evaluate the degree of inflation. For both the variant-specific differential and interaction analyses, DEseq2 results had notably inflated p-values (**SI Fig. 10E**). Given these findings, all downstream analyses were performed on results from mpralm.

#### Analysis of concordance and comparison to published MPRA studies

We first reviewed the overall number of tested CRS and the number of donors, replicates, scramble controls, and barcodes across 8 previously published MPRAs (**SI Data 2.18**). Our number of total replicates (4/condition; 2 donor, 2 technical replicates pooled across 6 wells) exceeds that of Kreimer et al. 2022 (1 donor, 3 reps)^110^, Rummel et al. 2023 (3 reps)^53^ and are comparable to the replicates in Deng et al. 2024 (5 replicates)^123^. Based on a minimum threshold of 10 barcodes per variant across all replicates, our mean number of barcodes (mean n=74 barcode) is comparable to the mean in Inoue et al. 2019 (n=70)^52^, Rummel et al. 2023 (n=45)^53^, and Deng et al. 2024 (n=64)^123^ and exceeds those in Tewhey et al. 2016 (n=20)^51^, Myint et al. 2019 (n=10)^78^, and Mulvey et al. 2021 (n=10)^122^. We specifically examined correlations between MPRA variant shifts in our study and in others testing psychiatric GWAS SNPs^54,78,122,123^ across different significance thresholds (**SI Data 2.17**). We compared activity and absolute overlap, reporting comparisons where there was enough overlap for Pearson’s correlation analysis (**SI Data 2.17**).

Of note, overlapping variants were not significantly correlated or there were too few to provide robust cross study validation. Many factors including the GWAS used, prioritization methods, size of the MPRA library, cell-type or environmental context, and the quantification methods (mpralm, MPRAnalyze, DESeq2, etc.) may explain this lack of overlap and correlation. However, this highlights a need to intentionally include a greater number of previously tested SNPs in MPRA library design and to replicate findings in the same cell-types and treatment paradigms independently.

#### Analysis of allelic shifts in MPRA activity and comparison to eQTL datasets

We first assessed the absolute overlap of all MPRA sequences post filtering and bulk adult PFC^75^, fetal brain^83^, and the CMC adult DLPFC eQTLs. For overlapping MPRA-emVars and eQTLs, we defined the percent concordant as the number of significant overlap SNPs with the same direction of effect in the MPRA-emVar and eQTL over the total number of overlapping SNPs. We further pared down this list by focusing on eQTL-eGene pairs that matched our prioritization by S-PrediXcan and coloc2. We calculated the Pearson’s Correlation Coefficient (r) between MPRA-emVar logFC and eQTL betas from post-mortem single-cell^84^ and bulk adult PFC^75^, and fetal brain^83^.

#### Variant-specific predicted binding affinity scoring with atSNP and MotifBreakR

To identify TF that may influence GxE interaction related to stress, performed we motif affinity testing for SNPs with significant allelic shifts as measured by lenti-MPRA (MPRA-emVars) using atSNP^154^. ENCODE derived TF motif binding PWM matrices were loaded and filtered for TF with expression in D24 iGLUTs, leaving 1573 motifs across 505 TFs. Genomic context (30bp half window size) of significant MPRA-emVars (p_FDR_<=0.1) for each condition were pulled with atSNP::LoadSNPData. SNP allele affinity scores were calculated for each motif with atSNP::ComputeMotifScore and p-values computed using atSNP::ComputePValues. Rank p-values were used to assess the significance of the SNP effect on the affinity change. Bonferroni multiple testing correction was performed along with calculation of Storey’s q-values and local FDR. Motif with significant SNP effects were tested for TF enrichments using hypergeometric tests with multiple testing correction across the total number of TFs tested. To identify TFs with binding patterns that predict variant-specific activity, SNP-motif pairs were filtered based on q_Storey_<0.05. Pearson’s Correlation Coefficients (r) were then calculated between MPRA-emVar activity (ref-vs-alt) and the logOdds that the reference allele enhanced affinity binding was assessed for each motif. To visualize allelic effects on predicted TF binding affinities for specific MPRA-emVars, we used the motifbreakR package^155^, filtered for strong allelic effects of TFs expressed in DIV24 iGLUTs.

#### Activity-by-Contact prediction of cue-specific regulatory element target genes

To predict target genes of condition-specific enhancer activity, we scored enhancer-gene interaction using STARE^86^. STARE combines an adapted Activity-By-Contact (ABC)^85^ interaction modeling with predicted TF binding affinities in regions to summarize these affinities at the gene level. For each condition, we filtered significant MPRA-emVar (p_FDR_ <0.1) or significant interaction MPRA-emVar (p_FDR_ <0.1) for overlap with matched cue-specific chromatin accessibility peaks (p_nom_<0.05) in iGLUTs. Enhancer “activity” for these variants was then calculated as the abs(logFC) compared to the alternative allele or baseline respectively, multiplied by the scaled peak AUC, thus incorporating crucial information about proximal cue-specific chromatin activity. Distal chromatin “contact” was measured from Hi-C data in iGLUTs^156^, which included one overlapping donor (2607). Genes were initially filtered by ABC score > 0.02 (default), but more stringent filtering was performed based on ABC score median thresholds prior to downstream enrichment analysis.

#### Over-representation analysis and biological theme comparison of correlated TFs and ABC genes

To identify pathway enrichments unique to cue-specific regulatory activity, we performed biological theme comparison using ClusterProfiler^153^ and gene set enrichment for GWAS catalogue risk genes using the GENE2FUNC query tool of FUMA GWAS^157^.

#### Enrichment analysis of convergence for risk loci using MAGMA

We tested ABC genes for enrichment with genetic risk of psychiatric, neurological, cardiometabolic, and immune disorders/traits [*Psychiatric*: attention-deficit/hyperactivity disorder (ADHD)^68^, anorexia nervosa (AN^158^, including binge-purge AN-BP and restrictive subtypes AN-R^159^), autism spectrum disorder (ASD)^3,160^, alcohol dependence (AUD)^161^, bipolar disorder (BIP, BIP-I, BIP-II)^70^ cannabis use disorder (CUD)^162^, major depressive disorder (MDD)^71^, obsessive-compulsive disorder (OCD)^72^, post-traumatic stress disorder (PTSD)^163^, and schizophrenia (SCZ)^1^, Cross Disorder (CxD)^164^, Tourette’s^165^, and neurotic personality traits^74^; *neurologic*: Alzheimer disease (AD)^67^, Parkinson disease (PD)^166^, amyotrophic lateral sclerosis (ALS)^167^, Multiple Sclerosis (MS)^168^, Epilepsy (Epi)^169^, migraine^170^, chronic pain^171^; *<u>Inflammatory-Gastrointestinal</u>*: Diverticular Disease (DivD)^172^, Gastro-esophageal reflux disease (GORD)^172^, Peptic Ulcer Disease (PUD)^172^, Inflammatory Bowel Disease (IBD)^172^, Irritable Bowel Syndrome (IBS)^172^; *<u>Cardiometabolic</u>*: Type-I Diabetes (T1D)^173^, Type-2 Diabetes (T2D)^174^, Metabolic Syndrome (MetS)^175^, Atrial Fibrillation (Afib)^176^, hypertension (HyperTen)^177^. *<u>Anthropometric:</u>* Body Mass Index (BMI)^178^, left-handedness (left hand)^179^ GWAS summary statistics] using multi-marker analysis of genomic annotation (MAGMA)^87^. SNPs were mapped to genes based on the corresponding build files for each GWAS summary dataset using the default method, snp-wise = mean (a test of the mean SNP association). A competitive gene set analysis was then used to test enrichment in genetic risk for a disorder across gene sets with an p_FDR_<0.05.

#### iGLUT neuron induction from non-clonal hiPSC-derived NPCs for phenotypic assays^56,57^

hiPSCs-derived NPCs were dissociated with Accutase Cell Detachment Solution (Innovative Cell Technologies, #AT-104), counted and transduced with rtTA (Addgene 20342) and *NGN2* (Addgene 99378) lentiviruses in StemFlex media containing 10 mM Thiazovivin (Millipore, #S1459). They were subsequently seeded at 1×10^6^ cells/well in the prepared 6-well plate. On day 1, medium was switched to non-viral induction medium (DMEM/F12 (Thermofisher, #10565018), 1% N-2 (Thermofisher, #17502048), 2% B-27-RA (Thermofisher, #12587010)) and doxycycline (dox) was added to each well at a final concentration of 1 μg/mL. At day 2, transduced hiPSCs were treated with 500 μg/mL G418 (Thermofisher, #10131035). At day 4, medium was replaced including 1 μg/mL dox and 4 μM cytosine arabinoside (Ara-C) to reduce the proliferation of non-neuronal cells. On day 5, immature NPC-iGLUTs were dissociated with Accutase Cell Detachment Solution (Innovative Cell Technologies, #AT-104), counted and seeded at a density of 1×10^6^ per well of a Matrigel-coated 12-well plate. Medium was switched to Brainphys neuron medium (Brainphys (STEMCELL, # 05790), 1% N-2, 2% B27-RA, 1 μg/mL Natural Mouse Laminin (Thermofisher, # 23017015), 10 ng/mL BDNF (R&D, #248), 10 ng/mL GDNF (R&D, #212), 500 μg/mL Dibutyryl cyclic-AMP (Sigma, #D0627), 200 nM L-ascorbic acid (Sigma, # A4403)). For seeding, 10 mM Thiazovivin (Millipore, #S1459), 500 μg/mL G418 and 4 μM Ara-C and 1 μg/mL dox were added. At day 6, medium was replaced with Brainphys neuron medium with 4 μM Ara-C and 1 μg/mL dox. Subsequently, 50% of the medium was replaced with fresh neuronal medium (lacking dox and Ara-C) once every other day until the NPC-iGLUTs were harvested at d21.

#### Neurite analysis

Day 7 NPC-iGLUTs were seeded as 1.5×10^4^ cells/well in a 96-well plate coated with 4x Matrigel at day 3 followed by half medium changes until the neurons were fixed at day 7. At day 5, NPC-iGLUTs were treated for 48hrs with either IL-6 (25ng/mL and 60 ng/mL), INFa-2b (100 IU/mL and 500 IU/mL), or matched vehicles. Following cue exposure, NPC-iGLUTs were fixed using 4% formaldehyde/sucrose in PBS with Ca^2+^ and Mg^2+^ for 10 minutes at room temperature (RT). Fixed cultures were washed twice in PBS and permeabilized and blocked using 0.1% Triton/2% Normal Donkey Serum (NDS) in PBS for two hours. Cultures were then incubated with primary antibody solution (1:1000 MAP2 anti chicken (Abcam, ab5392) in PBS with 2% NDS) overnight at 4°C. Cultures were then washed 3x with PBS and incubated with secondary antibody solution (1:500 donkey anti chicken Alexa 647 (Life technologies, A10042) in PBS with 2% NDS) for 1 hour at RT. Cultures were washed a further 3x with PBS with the second wash containing 1 μg/ml DAPI. Fixed cultures were then imaged on a CellInsight CX7 HCS Platform with a 20x objective (0.4 NA) and neurite tracing analysis performed using the neurite tracing module in the Thermo Scientific HCS Studio 4.0 Cell Analysis Software. 12 wells were imaged per condition across two control donor hiPSCs, with 9 images acquired per well for neurite tracing analysis. A one-way ANOVA with a post hoc Bonferroni multiple comparisons test was performed on data for neurite length per neuron using Graphpad Prism.

#### Synapse analyses

Commercially available primary human astrocytes (pHAs, Sciencell, #1800; isolated from fetal female brain) were seeded on D3 NPC-iGLUTs at 1.7×10^4^ cells per well on a 4x Matrigel-coated 96 W plate in neuronal media supplemented with 2% fetal bovine serum (FBS). NPC-iGLUTs were seeded over the astrocyte monolayer as 1.5×10^5^ cells/well at day 5 post induction. Half changes of neuronal media were performed twice a week until fixation. At day 13, NPC-iGLUTs + astrocyte co-cultures were treated with 200 nM Ara-C to reduce the proliferation of non-neuronal cells in the culture. At day 18, Ara-C was completely withdrawn by full medium change followed by half medium changes until the neurons were fixed at day 21. At day 21, NPC-iGLUTs + astrocytes co-cultures were treated for 48hrs with IL-6 (25ng/mL and 60 ng/mL), IFNα-2b (100 IU/mL and 500 IU/mL), or matched vehicles. Following exposure, NPC-iGLUTs were fixed and immune-stained as described previously, with an additional antibody stain for Synapsin1 (primary antibody: 1:500 Synapsin1 anti mouse (Synaptic Systems, 106 011); secondary antibody: donkey anti mouse Alexa 568 (Life technologies A10037)). Stained cultures were imaged and analyzed as above using the synaptogenesis module in the Thermo Scientific HCS Studio 4.0 Cell Analysis Software to determine SYN1+ puncta number, area, and intensity per neurite length in each image. 20 wells were imaged per condition across two control donor hiPSCs, with 9 images acquired per well for synaptic puncta analysis. A one-way ANOVA with a post hoc Bonferroni multiple comparisons test was performed on data for puncta number per neurite length using Graphpad Prism.

| Antibody | Species | Supplier | Product # | Dilution |
| --- | --- | --- | --- | --- |
| MAP2 | Ck | Abcam | ab5392 | 1:500 |
| SYNAPSIN1 | Ms | Synaptic Systems | 106 011 | 1:500 |
| Alexa 568 anti-Mouse | Ms | Life technologies | A10037 | 1:500 |
| Alexa 647 anti-Chicken | Ck | Life technologies | A10042 | 1:500 |

#### Multiple Electrode array (MEA)

Commercially available primary human astrocytes (pHAs, Sciencell, #1800; isolated from fetal female brain) were seeded on D3 NPC-iGLUTs at 1.7×10^4^ cells per well on a 4x Matrigel-coated 48 W MEA plate (catalog no. M768-tMEA-48W; Axion Biosystems) in neuronal media supplemented with 2% fetal bovine serum (FBS). At D5, iGLUTs were detached, spun down and seeded on the pHA cultures at 1.5×10^5^ cells per well. Half changes of neuronal media supplemented with 2% FBS were performed twice a week until day 42. At day 13, NPC-iGLUT + astrocyte co-cultures were treated with 200 nM Ara-C to reduce the proliferation of non-neuronal cells in the culture. At Day 18, Ara-C was completely withdrawn by full medium change. At day 26, NPC-iGLUTs + astrocytes co-cultures were treated for 48hrs with IL-6 (25 ng/mL and 60 ng/mL), IFNα-2b (100 IU/mL and 500 IU/mL), or matched vehicles. Following exposure, electrical activity of iGLUTs was recorded at 37°C using the Axion Maestro MEA reader (Axion Biosystems). Recording was performed via AxiS 2.4. Batch mode/statistic compiler tool was run following the final recording. Quantitative analysis of the recording was exported as Microsoft excel sheet. 6-12 wells were analyzed per condition across two control donor hiPSCs, with 16 electrodes per well for MEA. Data is log-transformed and normalized within each plate, prior to one-way ANOVA with a post hoc Bonferroni multiple comparisons test.

## STATEMENT OF ETHICS

Ethical approval was not required because the hiPSC lines, lacking association with any identifying information and widely accessible from a public repository, are thus not considered to be human subject research. Post-mortem brain data are similarly lacking identifiable information and are not considered human subject research.

## AUTHOR CONTRIBUTIONS

The paper was written by K.G.R., L.H. and K.J.B., with input from all authors. All high-throughput sequencing data and downstream analyses were performed by K.G.R.; L.D. provided code and guidance in conducting Bayesian colocalization analyses. S.L and M.J performed cue-specific ATAC experiments. S.E.W. performed ATAC-seq analysis. P.M.D performed dose-dependent morphological assays and analysis. M.F.G and K.G.R. performed cell-line clonalization. K.G.R performed MPRA library preparation and MPRA experiments with the assistance of M.F.G, S.C, S.C, and A.S.

## DATA AVAILABILITY

All source donor hiPSCs have been deposited at the Rutgers University Cell and DNA Repository (study 160; http://www.nimhstemcells.org/).

Full raw sequencing data will be made available through GEO upon publication. Summary statistics and accompanying code reported in this paper will be made available through Synapse (syn75962204; https://www.synapse.org/Synapse:syn75962204/wiki/)

## CODE AVAILABILITY

The full analysis pipeline (including code and processed data objects) used for analysis of RNA-seq, ATAC-seq, and MPRA data evaluation is publicly available through Synapse (syn75962204).

## GLOSSARY OF TERMS

MPRA sequences: All putative candidate regulatory sequences (CRS) from brain disease/trait GWAS prioritized for inclusion in the MPRA.
MPRA-CRSs: The subset of candidate regulatory sequences (CRS) with significant transcriptional activity resolved by MPRA (may be repressed or active compared to the mean transcriptional activity of the minimal reporter); MPRA-validated regulatory sequences.
MPRA-emVars: The subset of active MPRA (MPRA-active) CRS whereby single-nucleotide changes significantly altered transcriptional activity; MPRA-validated expression modulating variants.
Interaction MPRA-emVars: The subset of MPRA-emVars single-nucleotide changes that significantly alter transcriptional activity in a differential or dynamic manner between cues; MPRA-validated cue-by-variant expression modulating variants.

## REFERENCES

1 Trubetskoy, V. et al. Mapping genomic loci implicates genes and synaptic biology in schizophrenia. Nature 604, 502–508 (2022). 10.1038/s41586-022-04434-5

2 O’Connell, K. S. et al. Genomics yields biological and phenotypic insights into bipolar disorder. Nature 639, 968–975 (2025). 10.1038/s41586-024-08468-9

3 Grove, J. et al. Identification of common genetic risk variants for autism spectrum disorder. Nat Genet 51, 431–444 (2019). 10.1038/s41588-019-0344-8

4 Adams, M. J. et al. Trans-ancestry genome-wide study of depression identifies 697 associations implicating cell types and pharmacotherapies. Cell 188, 640–652.e649 (2025). 10.1016/j.cell.2024.12.002

5 Friligkou, E. et al. Gene discovery and biological insights into anxiety disorders from a large-scale multi-ancestry genome-wide association study. Nat Genet 56, 2036–2045 (2024). 10.1038/s41588-024-01908-2

6 Nievergelt, C. M. et al. Genome-wide association analyses identify 95 risk loci and provide insights into the neurobiology of post-traumatic stress disorder. Nat Genet 56, 792–808 (2024). 10.1038/s41588-024-01707-9

7 Kim, J. J. et al. Multi-ancestry genome-wide association meta-analysis of Parkinson’s disease. Nat Genet 56, 27–36 (2024). 10.1038/s41588-023-01584-8

8 Bellenguez, C. et al. New insights into the genetic etiology of Alzheimer’s disease and related dementias. Nat Genet 54, 412–436 (2022). 10.1038/s41588-022-01024-z

9 Uffelmann, E. et al. Genome-wide association studies. Nature Reviews Methods Primers 1, 59 (2021). 10.1038/s43586-021-00056-9

10 Abdellaoui, A., Yengo, L., Verweij, K. J. H. & Visscher, P. M. 15 years of GWAS discovery: Realizing the promise. Am J Hum Genet 110, 179–194 (2023). 10.1016/j.ajhg.2022.12.011

11 Cirnigliaro, M. et al. The contributions of rare inherited and polygenic risk to ASD in multiplex families. Proc Natl Acad Sci U S A 120, e2215632120 (2023). 10.1073/pnas.2215632120

12 Antaki, D. et al. A phenotypic spectrum of autism is attributable to the combined effects of rare variants, polygenic risk and sex. Nat Genet 54, 1284–1292 (2022). 10.1038/s41588-022-01064-5

13 Lipkin, W. I., Bresnahan, M. & Susser, E. Cohort-guided insights into gene-environment interactions in autism spectrum disorders. Nat Rev Neurol 19, 118–125 (2023). 10.1038/s41582-022-00764-0

14 Estes, M. L. & McAllister, A. K. Maternal immune activation: Implications for neuropsychiatric disorders. Science 353, 772–777 (2016). 10.1126/science.aag3194

15 Al-Haddad, B. J. S. et al. Long-term Risk of Neuropsychiatric Disease After Exposure to Infection In Utero. JAMA Psychiatry 76, 594–602 (2019). 10.1001/jamapsychiatry.2019.0029

16 Wu, Y., De Asis-Cruz, J. & Limperopoulos, C. Brain structural and functional outcomes in the offspring of women experiencing psychological distress during pregnancy. Mol Psychiatry 29, 2223–2240 (2024). 10.1038/s41380-024-02449-0

17 Han, V. X., Patel, S., Jones, H. F. & Dale, R. C. Maternal immune activation and neuroinflammation in human neurodevelopmental disorders. Nat Rev Neurol 17, 564–579 (2021). 10.1038/s41582-021-00530-8

18 Neuhaus, Z. F. et al. Maternal obesity and long-term neuropsychiatric morbidity of the offspring. Arch Gynecol Obstet 301, 143–149 (2020). 10.1007/s00404-020-05432-6

19 Kong, L., Nilsson, I. A. K., Brismar, K., Gissler, M. & Lavebratt, C. Associations of Different Types of Maternal Diabetes and Body Mass Index With Offspring Psychiatric Disorders. JAMA Netw Open 3, e1920787 (2020). 10.1001/jamanetworkopen.2019.20787

20 Smith, S. E., Li, J., Garbett, K., Mirnics, K. & Patterson, P. H. Maternal immune activation alters fetal brain development through interleukin-6. J Neurosci 27, 10695–10702 (2007). 10.1523/JNEUROSCI.2178-07.2007

21 Hsiao, E. Y., McBride, S. W., Chow, J., Mazmanian, S. K. & Patterson, P. H. Modeling an autism risk factor in mice leads to permanent immune dysregulation. Proc Natl Acad Sci U S A 109, 12776–12781 (2012). 10.1073/pnas.1202556109

22 Choi, G. B. et al. The maternal interleukin-17a pathway in mice promotes autism-like phenotypes in offspring. Science 351, 933–939 (2016). 10.1126/science.aad0314

23 Kim, E. et al. Maternal gut bacteria drive intestinal inflammation in offspring with neurodevelopmental disorders by altering the chromatin landscape of CD4(+) T cells. Immunity 55, 145–158 e147 (2022). 10.1016/j.immuni.2021.11.005

24 Garbett, K. A., Hsiao, E. Y., Kalman, S., Patterson, P. H. & Mirnics, K. Effects of maternal immune activation on gene expression patterns in the fetal brain. Translational psychiatry 2, e98 (2012). 10.1038/tp.2012.24

25 Gallagher, D. et al. Transient maternal IL-6 mediates long-lasting changes in neural stem cell pools by deregulating an endogenous self-renewal pathway. Cell stem cell 13, 564–576 (2013). 10.1016/j.stem.2013.10.002

26 Lombardo, M. V. et al. Maternal immune activation dysregulation of the fetal brain transcriptome and relevance to the pathophysiology of autism spectrum disorder. Mol Psychiatry 23, 1001–1013 (2018). 10.1038/mp.2017.15

27 Griego, E., Segura-Villalobos, D., Lamas, M. & Galvan, E. J. Maternal immune activation increases excitability via downregulation of A-type potassium channels and reduces dendritic complexity of hippocampal neurons of the offspring. Brain Behav Immun 105, 67–81 (2022). 10.1016/j.bbi.2022.07.005

28 Sarieva, K. et al. Pluripotent stem cell-derived neural progenitor cells can be used to model effects of IL-6 on human neurodevelopment. Dis Model Mech 16 (2023). 10.1242/dmm.050306

29 Sarieva, K. et al. Human brain organoid model of maternal immune activation identifies radial glia cells as selectively vulnerable. Mol Psychiatry 28, 5077–5089 (2023). 10.1038/s41380-023-01997-1

30 Goshi, N. et al. Direct effects of prolonged TNF-alpha and IL-6 exposure on neural activity in human iPSC-derived neuron-astrocyte co-cultures. Frontiers in cellular neuroscience 19, 1512591 (2025). 10.3389/fncel.2025.1512591

31 Yockey, L. J. et al. Type I interferons instigate fetal demise after Zika virus infection. Sci Immunol 3 (2018). 10.1126/sciimmunol.aao1680

32 Crow, Y. J. & Manel, N. Aicardi-Goutieres syndrome and the type I interferonopathies. Nat Rev Immunol 15, 429–440 (2015). 10.1038/nri3850

33 Fu, J. et al. Unraveling the regulatory mechanisms underlying tissue-dependent genetic variation of gene expression. PLoS genetics 8, e1002431 (2012). 10.1371/journal.pgen.1002431

34 GTEx Consortium, Battle, A., Brown, C. D., Engelhardt, B. E. & Montgomery, S. B. Genetic effects on gene expression across human tissues. Nature 550, 204–213 (2017). 10.1038/nature24277

35 Ota, M. et al. Dynamic landscape of immune cell-specific gene regulation in immune-mediated diseases. Cell 184, 3006–3021 e3017 (2021). 10.1016/j.cell.2021.03.056

36 Moore, S. R. et al. Sex differences in the genetic regulation of the blood transcriptome response to glucocorticoid receptor activation. Translational psychiatry 11, 632 (2021). 10.1038/s41398-021-01756-2

37 Yao, C. et al. Sex- and age-interacting eQTLs in human complex diseases. Hum Mol Genet 23, 1947–1956 (2014). 10.1093/hmg/ddt582

38 Linden, M. et al. Sex influences eQTL effects of SLE and Sjogren’s syndrome-associated genetic polymorphisms. Biol Sex Differ 8, 34 (2017). 10.1186/s13293-017-0153-7

39 Werling, D. M. et al. Whole-Genome and RNA Sequencing Reveal Variation and Transcriptomic Coordination in the Developing Human Prefrontal Cortex. Cell reports 31, 107489 (2020). 10.1016/j.celrep.2020.03.053

40 Cuomo, A. S. E. et al. Single-cell RNA-sequencing of differentiating iPS cells reveals dynamic genetic effects on gene expression. Nat Commun 11, 810 (2020). 10.1038/s41467-020-14457-z

41 Strober, B. J. et al. Dynamic genetic regulation of gene expression during cellular differentiation. Science 364, 1287–1290 (2019). 10.1126/science.aaw0040

42 Elorbany, R. et al. Single-cell sequencing reveals lineage-specific dynamic genetic regulation of gene expression during human cardiomyocyte differentiation. PLoS genetics 18, e1009666 (2022). 10.1371/journal.pgen.1009666

43 Seah, C. et al. Common genetic variation impacts stress response in the brain. bioRxiv, 2023.2012.2027.573459 (2023). 10.1101/2023.12.27.573459

44 Seah, C. et al. Modeling gene x environment interactions in PTSD using human neurons reveals diagnosis-specific glucocorticoid-induced gene expression. Nat Neurosci 25, 1434–1445 (2022). 10.1038/s41593-022-01161-y

45 Davenport, E. E. et al. Discovering in vivo cytokine-eQTL interactions from a lupus clinical trial. Genome biology 19, 168 (2018). 10.1186/s13059-018-1560-8

46 Hu, S. et al. Inflammation status modulates the effect of host genetic variation on intestinal gene expression in inflammatory bowel disease. Nat Commun 12, 1122 (2021). 10.1038/s41467-021-21458-z

47 Signer, R. et al. BMI-genome interactions regulate global gene expression with emphasis in brain and gut. Cell Genomics, 101280 (2026). 10.1016/j.xgen.2026.101280

48 Knowles, D. A. et al. Determining the genetic basis of anthracycline-cardiotoxicity by molecular response QTL mapping in induced cardiomyocytes. Elife 7 (2018). 10.7554/eLife.33480

49 Zhong, Y. et al. Leveraging drug perturbation to reveal genetic regulators of hepatic gene expression in African Americans. Am J Hum Genet 110, 58–70 (2023). 10.1016/j.ajhg.2022.12.005

50 Wolter, J. M. et al. Cellular Genome-wide Association Study Identifies Common Genetic Variation Influencing Lithium-Induced Neural Progenitor Proliferation. Biol Psychiatry 93, 8–17 (2023). 10.1016/j.biopsych.2022.08.014

51 Tewhey, R. et al. Direct Identification of Hundreds of Expression-Modulating Variants using a Multiplexed Reporter Assay. Cell 165, 1519–1529 (2016). 10.1016/j.cell.2016.04.027

52 Inoue, F., Kreimer, A., Ashuach, T., Ahituv, N. & Yosef, N. Identification and Massively Parallel Characterization of Regulatory Elements Driving Neural Induction. Cell stem cell 25, 713–727 e710 (2019). 10.1016/j.stem.2019.09.010

53 Rummel, C. K. et al. Massively parallel functional dissection of schizophrenia-associated noncoding genetic variants. Cell 186, 5165–5182 e5133 (2023). 10.1016/j.cell.2023.09.015

54 McAfee, J. C. et al. Systematic investigation of allelic regulatory activity of schizophrenia-associated common variants. Cell Genom 3, 100404 (2023). 10.1016/j.xgen.2023.100404

55 Lee, S. et al. Massively parallel reporter assay investigates shared genetic variants of eight psychiatric disorders. Cell 188, 1409–1424 e1421 (2025). 10.1016/j.cell.2024.12.022

56 Zhang, Y. et al. Rapid single-step induction of functional neurons from human pluripotent stem cells. Neuron 78, 785–798 (2013). 10.1016/j.neuron.2013.05.029

57 Ho, S. M. et al. Rapid Ngn2-induction of excitatory neurons from hiPSC-derived neural progenitor cells. Methods 101, 113–124 (2016). 10.1016/j.ymeth.2015.11.019

58 Nehme, R. et al. Combining NGN2 Programming with Developmental Patterning Generates Human Excitatory Neurons with NMDAR-Mediated Synaptic Transmission. Cell reports 23, 2509–2523 (2018). 10.1016/j.celrep.2018.04.066

59 Zheng, L. S. et al. Mechanisms for interferon-alpha-induced depression and neural stem cell dysfunction. Stem Cell Reports 3, 73–84 (2014). 10.1016/j.stemcr.2014.05.015

60 Rose-John, S., Jenkins, B. J., Garbers, C., Moll, J. M. & Scheller, J. Targeting IL-6 trans-signalling: past, present and future prospects. Nat Rev Immunol 23, 666–681 (2023). 10.1038/s41577-023-00856-y

61 Couch, A. C. M. et al. Acute IL-6 exposure triggers canonical IL6Ra signaling in hiPSC microglia, but not neural progenitor cells. Brain Behav Immun 110, 43–59 (2023). 10.1016/j.bbi.2023.02.007

62 Mzezewa, R. et al. A kainic acid-induced seizure model in human pluripotent stem cell-derived cortical neurons for studying the role of IL-6 in the functional activity. Stem cell research 60, 102665 (2022). 10.1016/j.scr.2022.102665

63 Kathuria, A., Lopez-Lengowski, K., Roffman, J. L. & Karmacharya, R. Distinct effects of interleukin-6 and interferon-gamma on differentiating human cortical neurons. Brain Behav Immun 103, 97–108 (2022). 10.1016/j.bbi.2022.04.007

64 Yvanka de Soysa, T., Therrien, M., Walker, A. C. & Stevens, B. Redefining microglia states: Lessons and limits of human and mouse models to study microglia states in neurodegenerative diseases. Semin Immunol 60, 101651 (2022). 10.1016/j.smim.2022.101651

65 Sofroniew, M. V. Astrocyte Reactivity: Subtypes, States, and Functions in CNS Innate Immunity. Trends Immunol 41, 758–770 (2020). 10.1016/j.it.2020.07.004

66 Laighneach, A., Desbonnet, L., Kelly, J. P., Donohoe, G. & Morris, D. W. Meta-Analysis of Brain Gene Expression Data from Mouse Model Studies of Maternal Immune Activation Using Poly(I:C). Genes (Basel) 12 (2021). 10.3390/genes12091363

67 Marioni, R. E. et al. GWAS on family history of Alzheimer’s disease. Translational psychiatry 8, 99 (2018). 10.1038/s41398-018-0150-6

68 Demontis, D. et al. Discovery of the first genome-wide significant risk loci for attention deficit/hyperactivity disorder. Nat Genet 51, 63–75 (2019). 10.1038/s41588-018-0269-7

69 Watson, H. J. et al. Genome-wide association study identifies eight risk loci and implicates metabo-psychiatric origins for anorexia nervosa. Nat Genet 51, 1207–1214 (2019). 10.1038/s41588-019-0439-2

70 Mullins, N. et al. Genome-wide association study of more than 40,000 bipolar disorder cases provides new insights into the underlying biology. Nat Genet 53, 817–829 (2021). 10.1038/s41588-021-00857-4

71 Howard, D. M. et al. Genome-wide meta-analysis of depression identifies 102 independent variants and highlights the importance of the prefrontal brain regions. Nat Neurosci 22, 343–352 (2019). 10.1038/s41593-018-0326-7

72 International Obsessive Compulsive Disorder Foundation Genetics, C. & Studies, O. C. D. C. G. A. Revealing the complex genetic architecture of obsessive-compulsive disorder using meta-analysis. Mol Psychiatry 23, 1181–1188 (2018). 10.1038/mp.2017.154

73 Huckins, L. M. et al. Analysis of Genetically Regulated Gene Expression Identifies a Prefrontal PTSD Gene, SNRNP35, Specific to Military Cohorts. Cell reports 31, 107716 (2020). 10.1016/j.celrep.2020.107716

74 Lo, M. T. et al. Genome-wide analyses for personality traits identify six genomic loci and show correlations with psychiatric disorders. Nat Genet 49, 152–156 (2017). 10.1038/ng.3736

75 Fromer, M. et al. Gene expression elucidates functional impact of polygenic risk for schizophrenia. Nat Neurosci 19, 1442–1453 (2016). 10.1038/nn.4399

76 Dobbyn, A. et al. Landscape of Conditional eQTL in Dorsolateral Prefrontal Cortex and Co-localization with Schizophrenia GWAS. Am J Hum Genet 102, 1169–1184 (2018). 10.1016/j.ajhg.2018.04.011

77 Barbeira, A. N. et al. Exploring the phenotypic consequences of tissue specific gene expression variation inferred from GWAS summary statistics. Nature Communications 9 (2018). 10.1038/s41467-018-03621-1

78 Myint, L. et al. A screen of 1,049 schizophrenia and 30 Alzheimer’s-associated variants for regulatory potential. Am J Med Genet B Neuropsychiatr Genet 183, 61–73 (2020). 10.1002/ajmg.b.32761

79 Ashuach, T. et al. MPRAnalyze: statistical framework for massively parallel reporter assays. Genome biology 20, 183 (2019). 10.1186/s13059-019-1787-z

80 Myint, L., Avramopoulos, D. G., Goff, L. A. & Hansen, K. D. Linear models enable powerful differential activity analysis in massively parallel reporter assays. BMC Genomics 20, 209 (2019). 10.1186/s12864-019-5556-x

81 Love, M. I., Huber, W. & Anders, S. Moderated estimation of fold change and dispersion for RNA-seq data with DESeq2. Genome biology 15, 550 (2014). 10.1186/s13059-014-0550-8

82 Abell, N. S. et al. Multiple causal variants underlie genetic associations in humans. Science 375, 1247–1254 (2022). 10.1126/science.abj5117

83 O’Brien, H. E. et al. Expression quantitative trait loci in the developing human brain and their enrichment in neuropsychiatric disorders. Genome biology 19, 194 (2018). 10.1186/s13059-018-1567-1

84 Bryois, J. et al. Cell-type-specific cis-eQTLs in eight human brain cell types identify novel risk genes for psychiatric and neurological disorders. Nat Neurosci 25, 1104–1112 (2022). 10.1038/s41593-022-01128-z

85 Fulco, C. P. et al. Activity-by-contact model of enhancer-promoter regulation from thousands of CRISPR perturbations. Nat Genet 51, 1664–1669 (2019). 10.1038/s41588-019-0538-0

86 Hecker, D., Behjati Ardakani, F., Karollus, A., Gagneur, J. & Schulz, M. H. The adapted Activity-By-Contact model for enhancer-gene assignment and its application to single-cell data. Bioinformatics 39 (2023). 10.1093/bioinformatics/btad062

87 de Leeuw, C. A., Mooij, J. M., Heskes, T. & Posthuma, D. MAGMA: generalized gene-set analysis of GWAS data. PLoS Comput Biol 11, e1004219 (2015). 10.1371/journal.pcbi.1004219

88 Liang, D. et al. Cell-type-specific effects of genetic variation on chromatin accessibility during human neuronal differentiation. Nat Neurosci 24, 941–953 (2021). 10.1038/s41593-021-00858-w

89 Degner, J. F. et al. DNase I sensitivity QTLs are a major determinant of human expression variation. Nature 482, 390–394 (2012). 10.1038/nature10808

90 Kaluscha, S. et al. Evidence that direct inhibition of transcription factor binding is the prevailing mode of gene and repeat repression by DNA methylation. Nat Genet 54, 1895–1906 (2022). 10.1038/s41588-022-01241-6

91 Grossman, S. R. et al. Systematic dissection of genomic features determining transcription factor binding and enhancer function. Proc Natl Acad Sci U S A 114, E1291–E1300 (2017). 10.1073/pnas.1621150114

92 Martinez-Corral, R. et al. Emergence of activation or repression in transcriptional control under a fixed molecular context. Proc Natl Acad Sci U S A 122, e2413715122 (2025). 10.1073/pnas.2413715122

93 Yockey, L. J. & Iwasaki, A. Interferons and Proinflammatory Cytokines in Pregnancy and Fetal Development. Immunity 49, 397–412 (2018). 10.1016/j.immuni.2018.07.017

94 Crow, Y. J. et al. Characterization of human disease phenotypes associated with mutations in TREX1, RNASEH2A, RNASEH2B, RNASEH2C, SAMHD1, ADAR, and IFIH1. Am J Med Genet A 167A, 296–312 (2015). 10.1002/ajmg.a.36887

95 Sullivan, K. D. et al. Trisomy 21 consistently activates the interferon response. Elife 5 (2016). 10.7554/eLife.16220

96 Escoubas, C. C. et al. Type-I-interferon-responsive microglia shape cortical development and behavior. Cell 187, 1936–1954 e1924 (2024). 10.1016/j.cell.2024.02.020

97 Kettwig, M. et al. Interferon-driven brain phenotype in a mouse model of RNaseT2 deficient leukoencephalopathy. Nat Commun 12, 6530 (2021). 10.1038/s41467-021-26880-x

98 Dumitriu, D. et al. Deciduous tooth biomarkers reveal atypical fetal inflammatory regulation in autism spectrum disorder. iScience 26, 106247 (2023). 10.1016/j.isci.2023.106247

99 Than, U. T. T. et al. Inflammatory mediators drive neuroinflammation in autism spectrum disorder and cerebral palsy. Scientific reports 13, 22587 (2023). 10.1038/s41598-023-49902-8

100 Mostafavi, S. et al. Type I interferon signaling genes in recurrent major depression: increased expression detected by whole-blood RNA sequencing. Mol Psychiatry 19, 1267–1274 (2014). 10.1038/mp.2013.161

101 Suzuki, K. et al. Microglial activation in young adults with autism spectrum disorder. JAMA Psychiatry 70, 49–58 (2013). 10.1001/jamapsychiatry.2013.272

102 Crow, Y. J. CNS disease associated with enhanced type I interferon signalling. The Lancet. Neurology 23, 1158–1168 (2024). 10.1016/S1474-4422(24)00263-1

103 Wamsley, B. et al. Molecular cascades and cell type-specific signatures in ASD revealed by single-cell genomics. Science 384, eadh2602 (2024). 10.1126/science.adh2602

104 Wu, W. L., Hsiao, E. Y., Yan, Z., Mazmanian, S. K. & Patterson, P. H. The placental interleukin-6 signaling controls fetal brain development and behavior. Brain Behav Immun 62, 11–23 (2017). 10.1016/j.bbi.2016.11.007

105 Rochfort, K. D. & Cummins, P. M. The blood-brain barrier endothelium: a target for pro-inflammatory cytokines. Biochem Soc Trans 43, 702–706 (2015). 10.1042/BST20140319

106 Braun, E. et al. Comprehensive cell atlas of the first-trimester developing human brain. Science 382, eadf1226 (2023). 10.1126/science.adf1226

107 Eze, U. C., Bhaduri, A., Haeussler, M., Nowakowski, T. J. & Kriegstein, A. R. Single-cell atlas of early human brain development highlights heterogeneity of human neuroepithelial cells and early radial glia. Nat Neurosci 24, 584–594 (2021). 10.1038/s41593-020-00794-1

108 Emani, P. S. et al. Single-cell genomics and regulatory networks for 388 human brains. Science 384, eadi5199 (2024). 10.1126/science.adi5199

109 Inoue, F. et al. A systematic comparison reveals substantial differences in chromosomal versus episomal encoding of enhancer activity. Genome Res 27, 38–52 (2017). 10.1101/gr.212092.116

110 Kreimer, A. et al. Massively parallel reporter perturbation assays uncover temporal regulatory architecture during neural differentiation. Nat Commun 13, 1504 (2022). 10.1038/s41467-022-28659-0

111 Yamamoto, R. et al. Tissue-specific impacts of aging and genetics on gene expression patterns in humans. Nat Commun 13, 5803 (2022). 10.1038/s41467-022-33509-0

112 Bryois, J. et al. Time-dependent genetic effects on gene expression implicate aging processes. Genome Res 27, 545–552 (2017). 10.1101/gr.207688.116

113 Huh, C. J. et al. Maintenance of age in human neurons generated by microRNA-based neuronal conversion of fibroblasts. Elife 5 (2016). 10.7554/eLife.18648

114 Mertens, J. et al. Directly Reprogrammed Human Neurons Retain Aging-Associated Transcriptomic Signatures and Reveal Age-Related Nucleocytoplasmic Defects. Cell stem cell 17, 705–718 (2015). 10.1016/j.stem.2015.09.001

115 Miller, J. D. et al. Human iPSC-based modeling of late-onset disease via progerin-induced aging. Cell stem cell 13, 691–705 (2013). 10.1016/j.stem.2013.11.006

116 Vera, E., Bosco, N. & Studer, L. Generating Late-Onset Human iPSC-Based Disease Models by Inducing Neuronal Age-Related Phenotypes through Telomerase Manipulation. Cell reports 17, 1184–1192 (2016). 10.1016/j.celrep.2016.09.062

117 Riessland, M. et al. Loss of SATB1 Induces p21-Dependent Cellular Senescence in Post-mitotic Dopaminergic Neurons. Cell stem cell 25, 514–530 e518 (2019). 10.1016/j.stem.2019.08.013

118 Ciceri, G. et al. An epigenetic barrier sets the timing of human neuronal maturation. Nature 626, 881–890 (2024). 10.1038/s41586-023-06984-8

119 Ernst, J. et al. Genome-scale high-resolution mapping of activating and repressive nucleotides in regulatory regions. Nat Biotechnol 34, 1180–1190 (2016). 10.1038/nbt.3678

120 Webber, C. Epistasis in Neuropsychiatric Disorders. Trends Genet 33, 256–265 (2017). 10.1016/j.tig.2017.01.009

121 Andreasen, N. C. et al. Statistical epistasis and progressive brain change in schizophrenia: an approach for examining the relationships between multiple genes. Mol Psychiatry 17, 1093–1102 (2012). 10.1038/mp.2011.108

122 Mulvey, B. & Dougherty, J. D. Transcriptional-regulatory convergence across functional MDD risk variants identified by massively parallel reporter assays. Translational psychiatry 11, 403 (2021). 10.1038/s41398-021-01493-6

123 Deng, C. et al. Massively parallel characterization of regulatory elements in the developing human cortex. Science 384, eadh0559 (2024). 10.1126/science.adh0559

124 Sheth, M. U. et al. Mapping enhancer-gene regulatory interactions from single-cell data. bioRxiv, 2024.2011.2023.624931 (2024). 10.1101/2024.11.23.624931

125 Gasperini, M. et al. A Genome-wide Framework for Mapping Gene Regulation via Cellular Genetic Screens. Cell 176, 377–390 e319 (2019). 10.1016/j.cell.2018.11.029

126 Ren, X. et al. High-throughput PRIME-editing screens identify functional DNA variants in the human genome. Mol Cell 83, 4633–4645 e4639 (2023). 10.1016/j.molcel.2023.11.021

127 Zhao, S. et al. A single-cell massively parallel reporter assay detects cell-type-specific gene regulation. Nat Genet 55, 346–354 (2023). 10.1038/s41588-022-01278-7

128 Wells, M. F. et al. Natural variation in gene expression and viral susceptibility revealed by neural progenitor cell villages. Cell stem cell 30, 312–332 e313 (2023). 10.1016/j.stem.2023.01.010

129 Gordon, M. G. et al. lentiMPRA and MPRAflow for high-throughput functional characterization of gene regulatory elements. Nat Protoc 15, 2387–2412 (2020). 10.1038/s41596-020-0333-5

130 Ghazi, A. R. et al. Design tools for MPRA experiments. Bioinformatics 34, 2682–2683 (2018). 10.1093/bioinformatics/bty150

131 Hunt, G. J., Freytag, S., Bahlo, M. & Gagnon-Bartsch, J. A. dtangle: accurate and robust cell type deconvolution. Bioinformatics 35, 2093–2099 (2019). 10.1093/bioinformatics/bty926

132 Dobin, A. et al. STAR: ultrafast universal RNA-seq aligner. Bioinformatics 29, 15–21 (2013). 10.1093/bioinformatics/bts635

133 Liao, Y., Smyth, G. K. & Shi, W. featureCounts: an efficient general purpose program for assigning sequence reads to genomic features. Bioinformatics 30, 923–930 (2014). 10.1093/bioinformatics/btt656

134 Leek, J. T., Johnson, W. E., Parker, H. S., Jaffe, A. E. & Storey, J. D. The sva package for removing batch effects and other unwanted variation in high-throughput experiments. Bioinformatics 28, 882–883 (2012). 10.1093/bioinformatics/bts034

135 Hoffman, G. E. & Schadt, E. E. variancePartition: interpreting drivers of variation in complex gene expression studies. BMC bioinformatics 17, 483 (2016). 10.1186/s12859-016-1323-z

136 Ritchie, M. E. et al. limma powers differential expression analyses for RNA-sequencing and microarray studies. Nucleic Acids Res 43, e47 (2015). 10.1093/nar/gkv007

137. 137 Wang, J. & Liao, Y. WebGestaltR: Gene Set Analysis Toolkit WebGestaltR. R package version 0.4.3., <https://CRAN.R-project.org/package=WebGestaltR> (2020).

138 Baines, K. J. et al. Maternal Immune Activation Alters Fetal Brain Development and Enhances Proliferation of Neural Precursor Cells in Rats. Front Immunol 11, 1145 (2020). 10.3389/fimmu.2020.01145

139 Dong, Y. et al. Transcriptomic profiling of the developing brain revealed cell-type and brain-region specificity in a mouse model of prenatal stress. BMC Genomics 24, 86 (2023). 10.1186/s12864-023-09186-8

140 Willer, C. J., Li, Y. & Abecasis, G. R. METAL: fast and efficient meta-analysis of genomewide association scans. Bioinformatics 26, 2190–2191 (2010). 10.1093/bioinformatics/btq340

141 r/TrimGalore: A wrapper around Cutadapt and FastQC to consistently apply adapter and quality trimming to FastQ files, with extra functionality for RRBS data (GitHub, 2023).

142. Andrews, S. FastQC A Quality Control tool for High Throughput Sequence Data. Babraham Bioinformatics

143 Ewels, P., Magnusson, M., Lundin, S. & Kaller, M. MultiQC: summarize analysis results for multiple tools and samples in a single report. Bioinformatics 32, 3047–3048 (2016). 10.1093/bioinformatics/btw354

144 Langmead, B. & Salzberg, S. L. Fast gapped-read alignment with Bowtie 2. Nat Methods 9, 357–359 (2012). 10.1038/nmeth.1923

145 Danecek, P. et al. Twelve years of SAMtools and BCFtools. GigaScience 10 (2021). 10.1093/gigascience/giab008

146 Zhang, Y. et al. Model-based analysis of ChIP-Seq (MACS). Genome biology 9, R137 (2008). 10.1186/gb-2008-9-9-r137

147 Li, Q., Brown, J. B., Huang, H. & Bickel, P. J. Measuring reproducibility of high-throughput experiments. The Annals of Applied Statistics 5, 1752–1779 (2011).

148 Hitz, B. C. et al. The ENCODE Uniform Analysis Pipelines. bioRxiv, 2023.2004.2004.535623 (2023). 10.1101/2023.04.04.535623

149 Robinson, J. T. et al. Integrative genomics viewer. Nat Biotechnol 29, 24–26 (2011). 10.1038/nbt.1754

150 DiffBind: differential binding analysis of ChIP-Seq peak data (Bioconductor, 2021).

151 Heinz, S. et al. Simple combinations of lineage-determining transcription factors prime cis-regulatory elements required for macrophage and B cell identities. Mol Cell 38, 576–589 (2010). 10.1016/j.molcel.2010.05.004

152 Yu, G., Wang, L. G. & He, Q. Y. ChIPseeker: an R/Bioconductor package for ChIP peak annotation, comparison and visualization. Bioinformatics 31, 2382–2383 (2015). 10.1093/bioinformatics/btv145

153 Yu, G., Wang, L. G., Han, Y. & He, Q. Y. clusterProfiler: an R package for comparing biological themes among gene clusters. OMICS 16, 284–287 (2012). 10.1089/omi.2011.0118

154 Zuo, C., Shin, S. & Keles, S. atSNP: Transcription factor binding affinity testing for regulatory SNP detection. Bioinformatics 31, 3353–3355 (2015).

155 Coetzee, S. G., Coetzee, G. A. & Hazelett, D. J. motifbreakR: an R/Bioconductor package for predicting variant effects at transcription factor binding sites. Bioinformatics 31, 3847–3849 (2015). 10.1093/bioinformatics/btv470

156 Rajarajan, P. et al. Neuron-specific signatures in the chromosomal connectome associated with schizophrenia risk. Science 362 (2018). 10.1126/science.aat4311

157 Watanabe, K., Taskesen, E., van Bochoven, A. & Posthuma, D. Functional mapping and annotation of genetic associations with FUMA. Nat Commun 8, 1826 (2017). 10.1038/s41467-017-01261-5

158 Duncan, L. et al. Significant Locus and Metabolic Genetic Correlations Revealed in Genome-Wide Association Study of Anorexia Nervosa. Am J Psychiatry 174, 850–858 (2017). 10.1176/appi.ajp.2017.16121402

159 Termorshuizen, J. D. et al. Genome-wide association studies of binge-eating behaviour and anorexia nervosa yield insights into the unique and shared biology of eating disorder phenotypes. medRxiv, 2025.2001.2031.25321397 (2025). 10.1101/2025.01.31.25321397

160 Matoba, N. et al. Common genetic risk variants identified in the SPARK cohort support DDHD2 as a candidate risk gene for autism. Translational psychiatry 10, 265 (2020). 10.1038/s41398-020-00953-9

161 Walters, R. K. et al. Transancestral GWAS of alcohol dependence reveals common genetic underpinnings with psychiatric disorders. Nat Neurosci 21, 1656–1669 (2018). 10.1038/s41593-018-0275-1

162 Johnson, E. C. et al. A large-scale genome-wide association study meta-analysis of cannabis use disorder. Lancet Psychiatry 7, 1032–1045 (2020). 10.1016/S2215-0366(20)30339-4

163 Nievergelt, C. M. et al. International meta-analysis of PTSD genome-wide association studies identifies sex- and ancestry-specific genetic risk loci. Nat Commun 10, 4558 (2019). 10.1038/s41467-019-12576-w

164 Cross-Disorder Group of the Psychiatric Genomics Consortium. Genomic Relationships, Novel Loci, and Pleiotropic Mechanisms across Eight Psychiatric Disorders. Cell 179, 1469–1482 e1411 (2019). 10.1016/j.cell.2019.11.020

165 Yu, D. et al. Interrogating the Genetic Determinants of Tourette’s Syndrome and Other Tic Disorders Through Genome-Wide Association Studies. Am J Psychiatry 176, 217–227 (2019). 10.1176/appi.ajp.2018.18070857

166 Nalls, M. A. et al. Identification of novel risk loci, causal insights, and heritable risk for Parkinson’s disease: a meta-analysis of genome-wide association studies. The Lancet. Neurology 18, 1091–1102 (2019). 10.1016/S1474-4422(19)30320-5

167 van Rheenen, W. et al. Common and rare variant association analyses in amyotrophic lateral sclerosis identify 15 risk loci with distinct genetic architectures and neuron-specific biology. Nat Genet 53, 1636–1648 (2021). 10.1038/s41588-021-00973-1

168 International Multiple Sclerosis Genetics, C. Multiple sclerosis genomic map implicates peripheral immune cells and microglia in susceptibility. Science 365 (2019). 10.1126/science.aav7188

169. International League Against Epilepsy Consortium on Complex, E. GWAS meta-analysis of over 29,000 people with epilepsy identifies 26 risk loci and subtype-specific genetic architecture. Nat Genet 55, 1471–1482 (2023). 10.1038/s41588-023-01485-w

170 Hautakangas, H. et al. Genome-wide analysis of 102,084 migraine cases identifies 123 risk loci and subtype-specific risk alleles. Nat Genet 54, 152–160 (2022). 10.1038/s41588-021-00990-0

171 Johnston, K. J. A. et al. Genome-wide association study of multisite chronic pain in UK Biobank. PLoS genetics 15, e1008164 (2019). 10.1371/journal.pgen.1008164

172 Wu, Y. et al. GWAS of peptic ulcer disease implicates Helicobacter pylori infection, other gastrointestinal disorders and depression. Nat Commun 12, 1146 (2021). 10.1038/s41467-021-21280-7

173 Chiou, J. et al. Interpreting type 1 diabetes risk with genetics and single-cell epigenomics. Nature 594, 398–402 (2021). 10.1038/s41586-021-03552-w

174 Mahajan, A. et al. Multi-ancestry genetic study of type 2 diabetes highlights the power of diverse populations for discovery and translation. Nat Genet 54, 560–572 (2022). 10.1038/s41588-022-01058-3

175 van Walree, E. S. et al. Disentangling Genetic Risks for Metabolic Syndrome. Diabetes 71, 2447–2457 (2022). 10.2337/db22-0478

176 Miyazawa, K. et al. Cross-ancestry genome-wide analysis of atrial fibrillation unveils disease biology and enables cardioembolic risk prediction. Nat Genet 55, 187–197 (2023). 10.1038/s41588-022-01284-9

177 Surendran, P. et al. Discovery of rare variants associated with blood pressure regulation through meta-analysis of 1.3 million individuals. Nat Genet 52, 1314–1332 (2020). 10.1038/s41588-020-00713-x

178 Speliotes, E. K. et al. Association analyses of 249,796 individuals reveal 18 new loci associated with body mass index. Nat Genet 42, 937–948 (2010). 10.1038/ng.686

179 Cuellar-Partida, G. et al. Genome-wide association study identifies 48 common genetic variants associated with handedness. Nat Hum Behav 5, 59–70 (2021). 10.1038/s41562-020-00956-y

