## Supplementary material for "Dynamic neuro-immune regulation of psychiatric risk loci in human neurons": SI Figures

**SUPPLEMENTAL DATA**

**Supplemental Data 1.** RNA-sequencing and ATAC-sequencing QC metrics, summary statistics, and gene set enrichments.

**Supplemental Data 2.** MPRA library SNP annotations, MPRA-emVar results

**SUPPLEMENTAL FIGURES**

**Supplemental Figure 1.** Experimental design for ATAC-seq, RNA-seq, and lenti-MPRA iGLUT experiments.

**Supplemental Figure 2.** IL-6 and IFNα receptor expression in mature iGLUT neurons.

**Supplemental Figure 3.** IL-6 and IFNα impact iGLUT gene expression, enriched for stress and immune-mediated signaling pathways.

**Supplemental Figure 4.** IL-6 and IFNα alter chromatin accessibility in mature iGLUTs.

**Supplemental Figure 5.** Correlations between cue-specific chromatin accessibility and gene expression changes in mature iGLUTs.

**Supplemental Figure 6.** Effect of IL-6 and IFNα on neurite outgrowth and synaptic puncta in iGLUTs.

**Supplemental Figure 7.** Selection of cis-expression quantitative trait loci (*cis*-eQTLs) and design of cross-disorder MPRA library.

**Supplemental Figure 8.** Correlations between MPRA biological and technical replicates.

**Supplemental Figure 9.** Comparison of CRS activity and variant-specific differential analysis results from MPRAnalyze, mpralm, and DEseq2.

**Supplemental Figure 10.** Comparison of cue-by-variant interaction analysis using MPRAnalyze, mpralm, and DEseq2.

**Supplemental Figure 11.** Scramble controls show significantly less transcriptional activity compared to eQTL and GWAS-prioritized sequences.

**Supplemental Figure 12.** The proportion of active CRS was significantly higher for eQTL and GWAS-prioritized sequences.

**Supplemental Figure 13.** Variant-specific effects were significantly correlated across conditions.

**Supplemental Figure 14**. After accounting for multi-mapping, MPRA-emVars concordant with bulk brain and single-cell excitatory eQTLs show cue-specificity but remain significantly correlated across conditions.

**Supplemental Figure 15.** Cue-response MPRA-emVars include GWAS SNPs associated with metabolic, cognitive, and affective traits.

**Supplemental Figure 16.** Variant-dependent TF binding affinities are differentially enriched based on cue exposure.

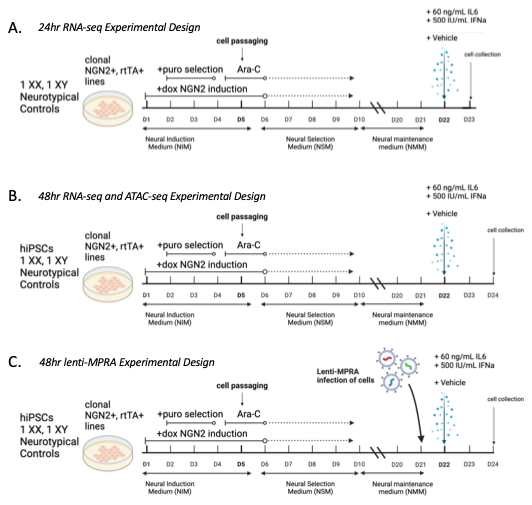
**Supplemental Figure 1. Experimental design for ATAC-seq, RNA-seq, and lenti-MPRA iGLUT experiments.** **(A)** Schematic for donor, maturity, and cue-matched 24hr RNA-sequencing experiments. **(B)** Schematic for donor, maturity, and cue-matched 48hr ATAC-sequencing and RNA-sequencing experiments. **(C)** Schematic for donor, maturity, and cue-matched lenti-MPRA experiments.

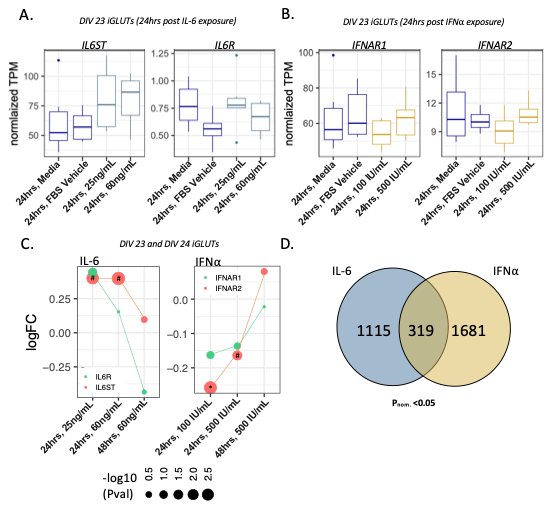

**Supplemental Figure 2.** **IL-6 and IFNα receptor expression in mature iGLUT neurons. (A)** Normalized basal expression levels of interferon-alpha, and interleukin-6 receptors in iGLUTs (DIV 23 or 24). Boxplots of normalized expression (TPM) of *IL6R* and *IL6ST* 24hr after exposure with either media, FBS vehicle, or IL-6 (25 ng/mL or 60 ng/mL) in 2 neurotypical donor lines with 3 technical replicates (n=6). **(B)** Boxplots of normalized expression (TPM) of *INFAR1* and *INFAR2* 24hr after exposure with either media, FBS vehicle, or IFNα (100 IU/mL or 500 IU/mL) in 2 neurotypical donor lines with 3 technical replicates (n=6). **(C)** LogFC of *IL6R*, *IL6ST*, *INFAR1*, *IFNΑR2* at 24hrs and 48hrs post exposure to IL-6 or IFNα compared to vehicle (#=nominal significant p-value <=0.05; *FDR significant).

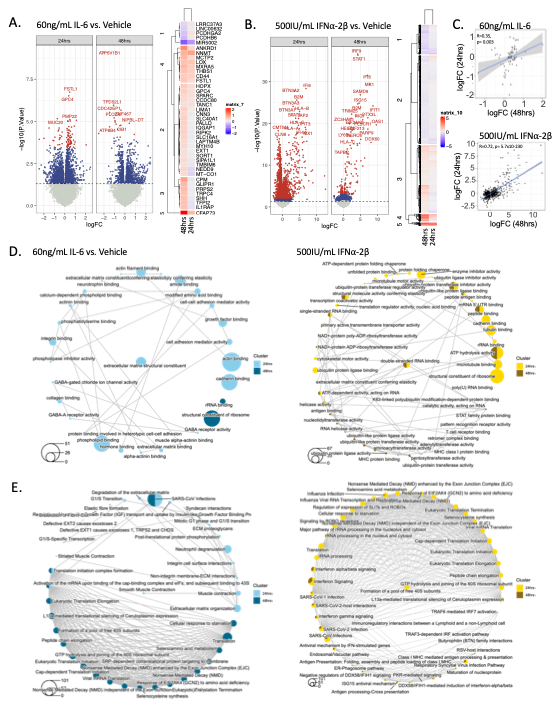
**Supplemental Figure 3. IL-6 and IFNα impact iGLUT gene expression, enriched for stress and immune-mediated signaling pathways.** An initial response to IL-6 and IFNα in mature iGLUTs was followed by partial return to homeostasis, highlighted by the greater overall number of differentially expressed genes (DEGs) seen in both conditions at 24hr compared to 48hr collection point. **(A)** DEGs at 24hrs and 48hrs post exposure to 60ng/mL IL-6 in mature iGLUTs (**SI Data 1.1-1.2**). **(B)** DEGs at 24hrs and 48hrs post exposure to 500 IU/mL IL-6 in mature iGLUTs (**SI Data 1.3-1.4**). **(C)** Significant DEGs were moderately and strongly correlated between the 24hr and 48hr collection points (IL-6: R=0.35, p=0.003; IFN-α: R=0.72, p=5.7x10^-230^), with evident shifts in the magnitude of effect. **(D)** Comparative Gene Ontology enrichments between 24hr and 28hr DEGs following exposure to IL-6 (teal) or IFNα (yellow). **(E)** Comparative REACTOME enrichments between 24hr and 28hr DEGs following exposure to IL-6 (teal) or IFNα (yellow).

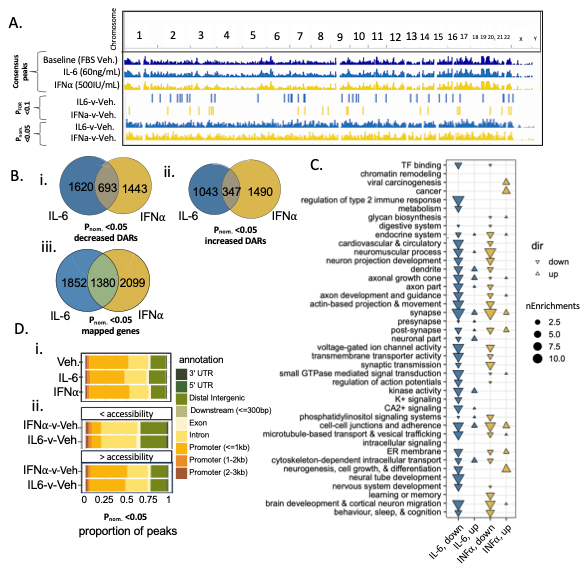
**Supplemental Figure 4. IL-6 and IFNα alter chromatin accessibility in mature iGLUTs. (A)** Consensus peaks and differentially accessible chromatin regions (DARs) across cue-exposure (**SI Data 1.8-1.11**), with the greatest number of differentially enriched peaks associated with IL-6 and IFNα (p_nom_<=0.05; p_FDR_<0.1) **(B)** Overlap of nominal DARs with increased (i) or decreased (ii) accessibility across conditions, and (iii) their mapped genes. **(C)** Absolute number of pathway enrichments (size of point) (p_FDR_<=0.05) by functional category across cue-responsive up and down regulated accessibility peaks. Color key: teal=IL-6, yellow= IFNα. **(D)** Proportion of chromatin accessibility peaks (p_nom_<=0.05) annotated as: 3’ UTR, 5’ UTRs, distal intergenic regions, downstream regions (<=300np), exons, introns, and promoters in relationship to the transcriptional state site (TSS) of mapped genes.

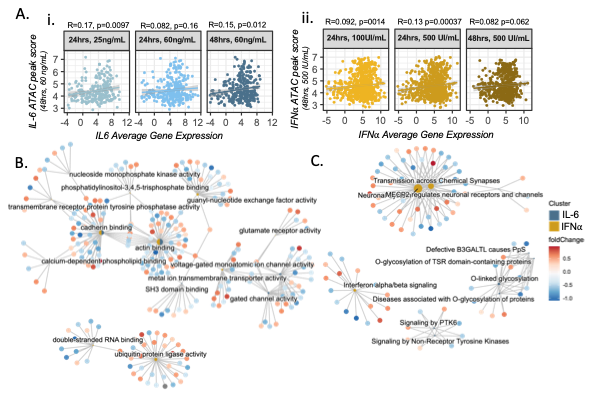

**Supplemental Figure 5. Correlations between cue-specific chromatin accessibility and gene expression changes in mature iGLUTs. (A)** Cue-responsive gene expression (average expression) and chromatin accessibility (ATAC peak score) were moderately correlated following 48hr IL-6 exposure (**A,i**: Pearson’s Correlation Coefficient (PCC); r=0.15, p=0.012) and 48hr, IFNα exposure (**A,ii**: r=0.082, p=0.062). **(B-C)** Enrichments of differentially expressed genes (DEGs) overlapping with chromatin accessibly peaks (DARs). Comparative enrichment analysis revealed IFNα specific enrichments (e.g., Interferon-alpha signaling, neuronal synaptic transmission, double-stranded RNA binding, and ubiquitin protein modification) and IL-6-specific enrichments (e.g., O-linked glycosylation, signaling by PTK6, MECP2 regulation or neuronal receptors, calcium-dependent phospholipid binding and SH3 binding), and shared enrichments in actin and cadherin binding (**SI Data 1.11**).

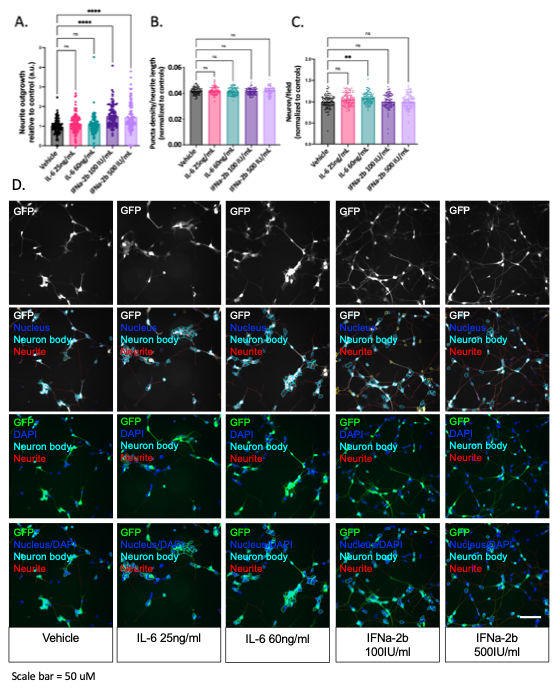
**Supplemental Figure 6.** **Effect of IL-6 and IFNα on neurite outgrowth and synaptic puncta in iGLUTs.** **(A)** Exposure to 100 IU/mL and 500 IU/mL IFNα-2b but not 25ng/ml or 60ng/mL IL-6, increased neurite outgrowth (D7) relative to vehicle control conditions. **(B)** Synaptic puncta density (relative expression of Synapsin 1 (SYN1) in D21 neurons co-cultured with astrocytes) following cue-exposure. The relative number of neurons compared was not significantly less compared to controls, suggesting that application of IL-6 and IFNα at the doses did not result in significant cell death. **(C)** Proportion of neurons following exposure to IL-6 (60ng/mL). N = minimum of 2 independent experiments across 2 donor lines with 12 technical replicates per condition and 9 images analyzed per replicate. One way ANOVA with post-hoc Bonferroni multiple comparisons test * = P_bon_<0.05; ** = P_bon_<0.01; *** = P_bon_<0.001; **** = P_bon_<0.0001. **(D)** Representative traces of neuron body across conditions and doses. Scale bar: 50µm.

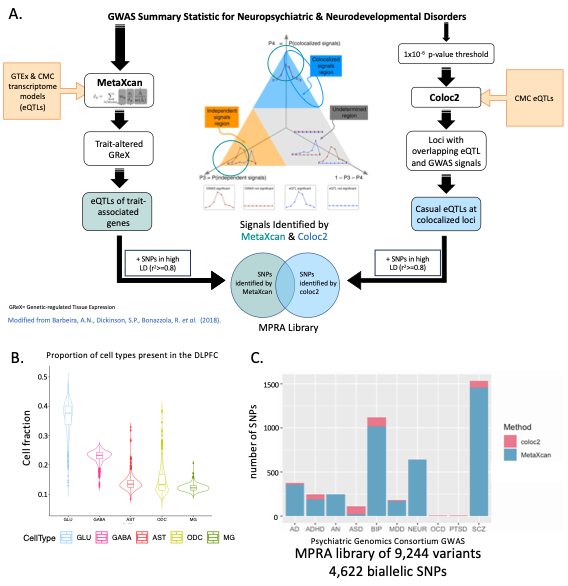

**Supplemental Figure 7.** **Selection of cis-expression quantitative trait loci (*cis*-eQTLs) and design of cross-disorder MPRA library. (A)** Variant selection from ten GWAS (AD, ADHD, AN, ASD, BIP, MDD, OCD, PTSD, SCZ, and NEU**; SI Data 2.1**) applied two selection approaches: (1) Bayesian co-localization (using coloc2^130,131^) and (2) transcriptomic imputation (S-PrediXcan^132,133^) (**Fig. 6**). For method 1, significant GWAS loci were identified as those LD r^2^ > 0.1 to lead associated SNPs (p<1x10^-6^). Colocalization between GWAS SNPs and CMC DLPFC cis-eQTLs (FDR < 0.05) for overlapping genes was tested using coloc2^131^; loci with PPH4 >= 0.5 were considered moderately to strongly co-localized. The most probable causal eQTLs from these loci were selected, along with all SNPs in high LD (r^2^ >= 0.8) (**SI Data 2.2-2.3**). For method 2, trait-associated CMC DLPFC S-PrediXcan genes [p<~4.64x10^-6^ (0.05/10786)] were selected for AD, ADHD, AN, ASD, BIP, MDD, OCD, PTSD, SCZ, and NEU. All SNPs within the predictor models of significant genes (3-15/gene) and all SNPs in high LD (r^2^ >= 0.8) with them were selected (**SI Data 2.4-2.5**). The following sets of benchmark variants were included: 100 active and 100 inactive CRSs were selected from SNPs with the greatest and least transcriptional shifts based on a previous SCZ and AD MPRA^115^ that were present in the CMC DLPFC dataset (**SI Data 2.7**). Coloc2 negative controls were selected from significant BIP GWAS loci that did not colocalize (PPH3 > 0.9) and were not significant CMC DLPFC eQTLs. 100 scramble sequences were used as controls, representing activity of the minimal promoter. (**B)** Estimate proportion of cell types present in the CMC postmortem dorsolateral prefrontal cortex (DLPFC) by cell-type deconvolution. **(C)** Proportion of test SNPs by prioritization method and GWAS summary statistics.

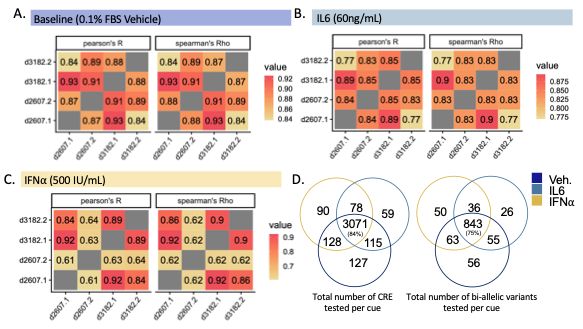
**Supplemental Figure 8.** **Correlations between MPRA biological and technical replicates. (A-C)** Transcriptional activity, measured as the normalized log2 RNA/DNA ratios, is strongly correlated between replicates at baseline (**A**), with IL-6 (**B**) and IFNα (**C)**: Pearson’s R=0.61-0.93, mean=82.5; Spearman’s Rho=0.62-0.93, mean=82.5. **(D)** Following removal of low-DNA count sequences, 3440 (Baseline), 3322 (IL-6), and 3366 (IFNα) CRSs were captured with a minimum requirement of 10 barcodes across replicates (mean 45 barcodes/CRS). Of these, 1018 (Base), 960 (IL-6), and 992 (IFNα), were biallelic and used in downstream analysis of variant-specific effects. The number of barcodes per CRS were highly correlated between shared across all conditions (Pearson’s R=0.997-0.999) (**SI Data 2.11**).

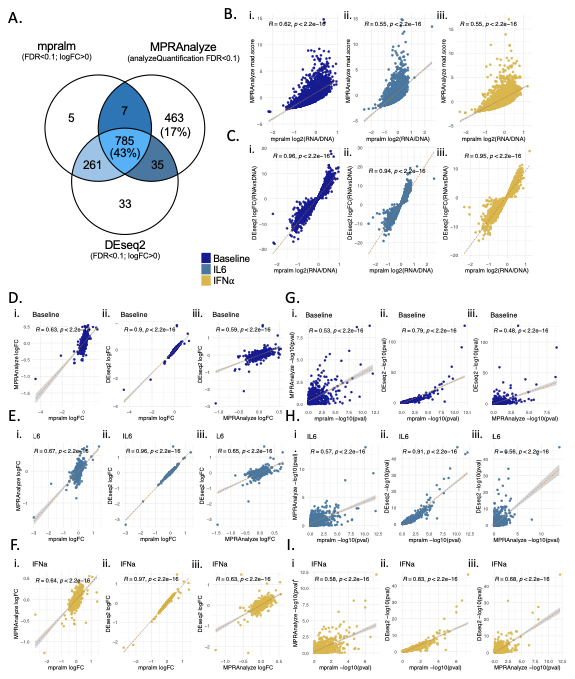

**Supplemental Figure 9.** **Comparison of CRS activity and variant-specific differential analysis results from MPRAnalyze, mpralm, and DEseq2.** Quantification of CRS activity and variant-specific differential analysis was performed using three previously published approaches: MPRAnalyze, mpralm, and DEseq2. **(A)** At baseline, 43% of active CRS (p_FDR_<=0.1) were detected in all methods. mpralm and DEseq2 identified the same 1046 CRS, with only 12 additional CRS identified by mpralm and not DEseq2. **B-C.** Estimated change in CRS activity compared to the mean activity of all CRS (i.e. shift up or down compared to the activity of the minimal promoter). Activity scores were **(B)** moderately correlated between MPRAnalyze (y axis) and mpralm (x axis) (Pearson’s correlation coefficient, R=0.55-0.62) and **(C)** highly correlated between DEseq2 (y.axis) and mpralm (x.axis) (R=0.94-0.96). **(D-I)** Variant-specific differential analyses between methods across contexts: baseline (**D**), IL-6 (**E**) and IFNα (**F):** logFC of variant-effects were moderately correlated between MPRAnalyze and mpralm (r=0.63-0.67) **(i)** and MPRAnalyze and DEseq2 (R=0.59-0.65) **(iii)** and strongly correlated between mpralm and DESeq2 (R=0.9-0.97) **(ii).** Likewise, -log10(pvalue) correlations **(G-I)** were strongest between **(ii)** mpralm and DEseq2 (R=0.79-0.91) and moderate between MPRAnalyze and **(i)** mpralm (R=0.53-0.58) or **(ii)** DEseq2 (R=0.48-0.68).

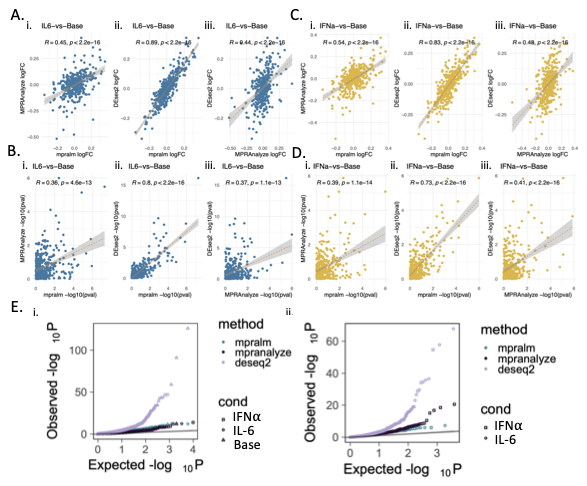
**Supplemental Figure 10.** **Comparison of cue-by-variant interaction analysis from MPRAnalyze, mpralm, and DEseq2.** Cue-by-variant interaction tests were compared across methods. As in the quantification and variant-specific differential analysis (**SI. Fig. 10**), mpralm and DESeq2 **(A-B)** logFC and **(C-D)** -log10(p-values) values were strongly correlated (logFC Pearson’s R=0.83-0.89, -log10(p-value) R=0.73-0.8) while MPRAnalyze results moderately correlated with either method. Overall, mpralm and DEseq2 results shared the highest concordance. **(E)** Comparisons of observed and expected p-values for the variant-specific differential analysis **(i)** and the cue-by-variant interaction test (ii) showed noticeable p-value inflation in DEseq2 analyses.

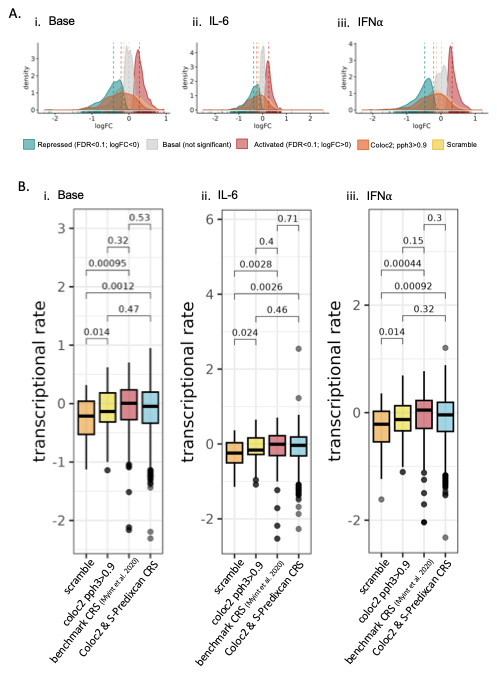
**Supplemental Figure 11. Scramble controls show significantly less transcriptional activity compared to eQTL and GWAS-prioritized CRS.** **(A)** Distribution plots of -logFC values (difference in the activity of a CRS when compared to the average rate of transcription by the minimal promoter). Based on the mpralm quantification analysis, we define three groups (a) activated (red; p_FDR_<0.1, logFC>0), (b) repressed (teal; p_FDR_<0.1, logFC<0), and (c) basal (grey; not significant). The distribution of activity in scramble controls and negative colocalization controls (pph3>0.9) overlap more with basal (not significant) CRS as expected. **(B)** The activity of scramble controls was significantly less than active elements from Myint et al. 2020 (Students T-Test Baseline: p-value=0.00092, IL-6 p=0.0028, and IFNα=0.00044), coloc2-negative controls (SNPs from non-colocalized loci, pph3>0.9; Students T-Test Baseline: p-value=0.014, IL-6 p=0.024, and IFNα=0.014), and our eQTL and GWAS-prioritized CRS (Students T-Test Baseline: p-value=0.0012, IL-6 p=0.0026, and IFNα=0.00092). Boxplots of transcriptional rate of scramble controls, negative controls based on colocalization, empirical controls, and MPRA inserts across conditions with lower and upper hinges (the 25th and 75th percentiles), lower and upper whisker that extends from the hinge to the largest or smallest value no further than 1.5 * IQR (inter-quartile range). Outlying points are plotted individually. Corresponds to analyses in **Figure 1.**

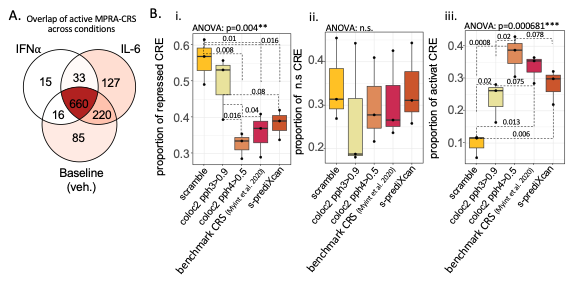
**Supplemental Figure 12. The proportion of active CRS was significantly higher for eQTL and GWAS-prioritized sequences (A)** Overlap of active MPRA-CRS by condition corresponding to **Figure 1**. **(B)** Boxplots with each experiment treated as a replicate, comparing the overall proportion of **(i)** repressed (mpralm p_FDR_<=0.1, logFC<0), **(ii)** not significant (mpralm p_FDR_>0.1), and **(iii)** activated (mpralm p_FDR_<=0.1, logFC>0). Proportions of repressed (one-way paired ANOVA p=0.004, post-hoc Student’s paired t-test) and active CRS were significantly different across prioritization methods (one-way paired ANOVA p=0.000681, post-hoc paired Student’s t-test). Scramble controls had the highest proportion of repressed and lowest proportion of active sequences, followed by negative controls selected from non-colocalized loci (coloc2, pph3>0.9). Among prioritized CRSs, those selected from colocalization had the highest proportion of active regions, followed by GWAS-based benchmark CRSs, and then s-PrediXcan-based CRSs (trending p=0.078). Corresponds to analyses in **Figure 1.** Boxplots with lower and upper hinges (the 25th and 75th percentiles), lower and upper whisker that extends from the hinge to the largest or smallest value no further than 1.5 * IQR (inter-quartile range). Individual samples (unique MPRA experiments n=3) are represented by individual points. Differences in the proportions was evaluated by two-way ANOVA followed by paired two-sided T-tests after testing for assumptions of normalcy (Shapiro-Wilk’s Test) and equal variance (Levene’s Test).

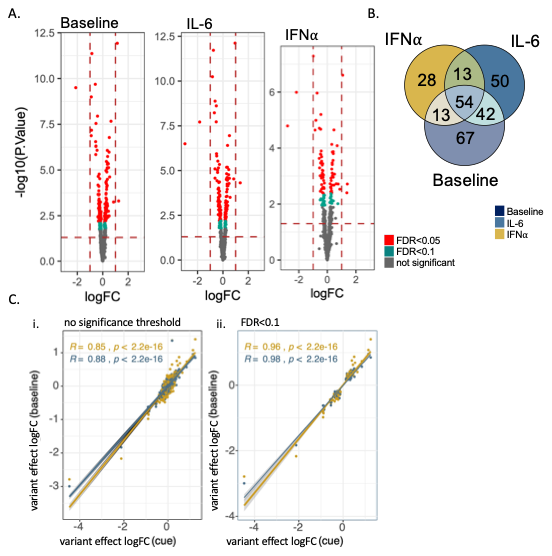

**Supplemental Figure 13. Variant-specific effects were significantly correlated across conditions.** 31% of baseline MPRA-emVars were significant after exposure to IL-6 and IFNα. **(A)** Volcano plots of differentially active variants (MPRA-emVars) at baseline and following exposure to IL-6 (60ng/mL), and IFNα (500UI). QTLs at p_FDR_<=0.1 (teal) or p_FDR_<=0.05 (terracotta). **(B)** Overlap of significant MPRA-emVars (p_FDR_<=0.1) across conditions. **(C)** Correlation coefficients between MPRA-emVar effects at baseline (y-axis) and cue exposure (x-axis) across two thresholds: **(i)** no significance threshold; **(ii)** p_FDR_<=0.1. Corresponds to analyses in **Fig. 1-3.**

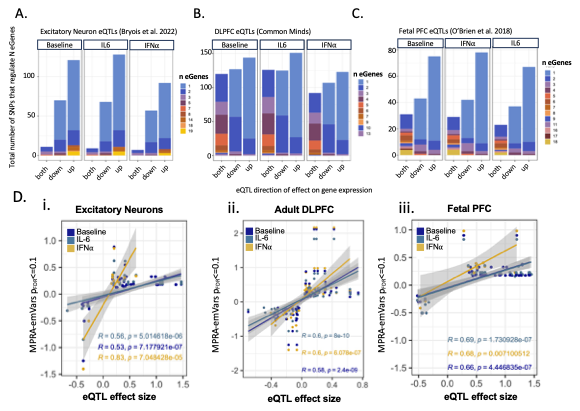

**Supplemental Figure 14**. **After accounting for multi-mapping, MPRA-emVars concordant with bulk brain and single-cell excitatory eQTLs show cue-specificity, while remaining significantly correlated across conditions.** In eQTL analysis, a single SNP can associate with multiple eGenes, which may have variable magnitude and directions of effect. **(A-C)** To contextualize our results in **Fig 2,** whether SNPs regulate all gene targets in the same direction (all up-regulation or all down-regulation) or with opposing effects on gene expression based (both) is considered across the total number of SNPs that significantly regulate one or multiple eGenes using **(A)** single-cell excitatory neuron, **(B)** bulk adult DLPFC, and **(C)** fetal PFC eQTLs is presented. **(D)** Correlations between MPRA and eQTLs. The x-axis displays Pearson’s correlations between concordant MPRA-emVar and **(i)** adult excitatory neuron eQTLs, **(ii)** adult DLPFC eQTLs, and **(iii)** fetal PFC eQTLs at a p_FDR_<=0.1 significance threshold (dark blue=baseline, light blue=IL-6, gold=IFNα). Corresponds to analyses in **Fig. 2**.

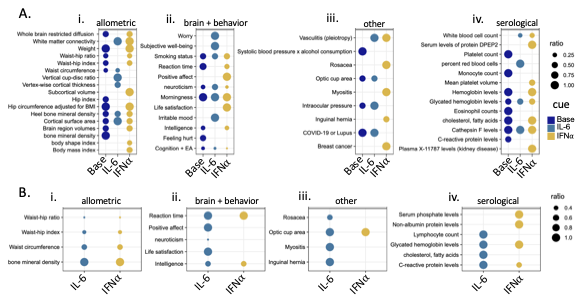

**Supplemental Figure 15. Cue-response MPRA-emVars include GWAS SNPs associated with metabolic, cognitive, and affective traits.** GWAS annotations of **(A)** MPRA-emVars and **(B)** interaction MPRA-emVars indicate cue-specific regulation of psych-psych pleiotropy or psych-cardiometabolic pleiotropy (**Fig. 3**, GWAS Catalogue; **SI Data 2.19**), highlighting allometric **(i),** behavioral **(ii)** and serological (**iv**) measures**.** Size of the points indicates the proportion of MPRA-emVars compared to the absolute number of overlapping MPRA and GWAS SNPs. Corresponds to analyses in **Figure 3**.

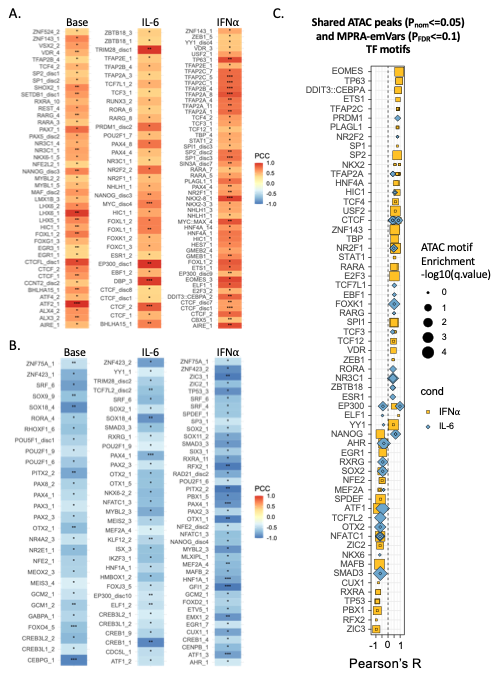
**Supplemental Figure 16. Variant-dependent TF binding affinities are differentially enriched based on cue exposure.** Allele-specific effects on transcription factor (TF) binding affinities were tested across 1572 binding motifs across 505 TFs expressed in mature iGLUTs. **(A-B)** TF motif-SNP interactions with predicted difference in binding based on variant (q_Storey_<0.05) were compared with measured allelic effects across MPRA-emVars (p_FDR_<=0.1). Top TF motifs both **(A)** positively and **(B)** negatively correlated with enhancer activity were largely unique across conditions. Heatmap of Pearson’s correlation coefficients (PCC) of TF motifs whose predicted effect on activity corelates with MPRA-emVar allelic effects (**SI Data 2.20**; p<0.08^#^, p<0.05*, p<0.01**, p<0.001***). **(C)** Many of the top predicted MPRA-emVar regulating transcription factors overlapped with motif enrichments based on chromatin accessibility in donor matched mature iGLUTs exposed to vehicle, IL-6, or IFNα (cue-specific ATAC peaks). Corresponds to analyses in **Figure 4**.
